# Fast and inexpensive whole genome sequencing library preparation from intact yeast cells

**DOI:** 10.1101/2020.09.03.280990

**Authors:** Sibylle C Vonesch, Shengdi Li, Chelsea Szu Tu, Bianca P Hennig, Nikolay Dobrev, Lars M Steinmetz

## Abstract

Through the increase in the capacity of sequencing machines massively parallel sequencing of thousands of samples in a single run is now possible. With the improved throughput and resulting drop in the price of sequencing, the cost and time for preparation of sequencing libraries have become the major bottleneck in large-scale experiments. Methods using a hyperactive variant of the Tn5 transposase efficiently generate libraries starting from cDNA or genomic DNA in a few hours and are highly scalable. For genome sequencing, however, the time and effort spent on genomic DNA isolation limits the practicability of sequencing large numbers of samples. Here, we describe a highly scalable method for preparing high quality whole-genome sequencing libraries directly from yeast cultures in less than three hours at 34 cents per sample. We skip the rate-limiting step of genomic DNA extraction by directly tagmenting yeast spheroplasts and add a nucleosome release step prior to enrichment PCR to improve the evenness of genomic coverage. Resulting libraries do not show any GC-bias and are comparable in quality to libraries processed from genomic DNA with a commercially available Tn5-based kit. We use our protocol to investigate CRISPR/Cas9 on- and off-target edits and reliably detect edited variants and shared polymorphisms between strains. Our protocol enables rapid preparation of unbiased and high-quality, sequencing-ready indexed libraries for hundreds of yeast strains in a single day at a low price. By adjusting individual steps of our workflow we expect that our protocol can be adapted to other organisms.

## BACKGROUND

Whole-genome sequencing is a powerful tool in genomics research by providing an unbiased and comprehensive view of the genetic alterations present in a cell. Genomic information is instrumental for identifying mutations that underlie observed phenotypes and for pinpointing any collateral damage that can occur as a side product of mutagenesis. As sequencing costs continue to drop, the cost and time for preparation of sequencing libraries have become the major limiting factor for large-scale genome sequencing experiments.

The most rapid and scalable library preparation methods use a hyperactive variant of the Tn5 transposase that fragments double-stranded DNA and ligates synthetic oligonucleotide adapters required for Illumina sequencing in a 5 minute reaction^1^ (Illumina). While the one-step tagmentation reaction greatly simplifies library preparation workflows compared to traditional, multi-step methods, and scales to the parallel processing of hundreds of samples, the cost of commercial reagents prevents its use in large-scale projects for most laboratories. We^2^ and others^3^ have previously described a robust Tn5 transposase purification strategy and accompanying library preparation protocol that allows generating sequencing libraries with comparable quality but at dramatically reduced cost compared to commercial solutions. In addition to a hyperactive Tn5 enzyme variant carrying the previously reported missense mutations E54K^4,5^ and L372P^6^, which increase the DNA-binding efficiency and reduce inhibitory effects on Tn5 activity, respectively, we introduced a second Tn5 construct carrying an additional amino acid substitution (R27S) in the DNA-binding domain^2^, which allows adjusting the fragment size distribution based on enzyme concentration during tagmentation.

With Tn5-based adapter insertion using homemade enzymes, genomic DNA isolation becomes the major bottleneck limiting the practicability of sequencing large numbers of genomes. Analogous to colony PCR, where lysed bacterial cells or yeast spheroplasts are added to a PCR reaction without prior genomic DNA isolation, we hypothesized that yeast whole-genome sequencing library preparation could be simplified by applying tagmentation directly to cells. A similar strategy incorporating heat-based lysis prior to tagmentation was successfully applied for whole-genome^1^ and plasmid^7^ sequencing of bacterial cells. In contrast to bacteria, the yeast *S. cerevisiae* has a rigid cell wall that prevents efficient heat-based lysis. To facilitate cellular lysis the lytic enzyme Zymolyase^8^, which actively degrades the cell walls of various yeasts, thereby exposing the fragile spheroplast, is routinely used in genomic DNA extraction^9,10^ and colony PCR^11^ protocols. With an MW of 138 kDA the active Tn5 dimer-adapter complex does not far exceed the size of average polymerase enzymes (~90kDa), such that treatment with Zymolyase should provide sufficient access to genomic DNA to Tn5. Furthermore, a similar method is used for Tn5-based adapter insertion into accessible DNA in yeast ATAC-seq^12^ protocols.

Here, we report a simplified whole-genome sequencing library preparation protocol. By skipping a conventional genomic DNA isolation step our method enables preparing sequencing-ready libraries directly from overnight yeast cultures in less than three hours at a cost of 34 cents per sample, while scaling to the parallel processing of hundreds of samples. In place of a lengthy genomic DNA isolation step, we incubate saturated yeast cultures with Zymolyase and apply tagmentation directly to the resulting spheroplasts. A nucleosome release step prior to enrichment PCR improves the evenness of genomic coverage. Resulting libraries are comparable in quality to libraries processed from extracted genomic DNA with a commercially available Tn5-based kit (Nextera XT, Illumina). We demonstrate the simplicity and unbiased nature of the method by investigating CRISPR/Cas9 on- and off-target editing outcomes in a panel of yeast strains, reliably detecting edited variants and shared polymorphisms between strains. Direct preparation of sequencing libraries from yeast cultures without DNA isolation enables massively scaled genome sequencing experiments that have so far been hampered by the time and effort spent on genomic DNA isolation. At 34 cents library preparation costs are approximately 5-10-fold lower compared to libraries prepared from gDNA extracted using the most simple and scalable kit-based methods. By adjusting steps like initial lysis to specific sample requirements we anticipate that our protocol can be applied to other microbial cells, mammalian cells or tissues.

## RESULTS

### Initial comparison of libraries prepared from genomic DNA versus intact cells

We hypothesized that tagmentation could be applied directly to yeast spheroplasts to simplify whole-genome sequencing library preparation by skipping genomic DNA isolation. Figure 1 illustrates the extraction-free library preparation workflow using homemade Tn5_R27S,E54K,L372P_. Instead of extracting genomic DNA, cells from saturated overnight cultures are exposed to Zymolyase treatment to digest the cell wall, and the Tn5 adapter complex is added directly to the spheroplasts (Fig. 1). As nucleosomes could provide a barrier for polymerase during library amplification we compared two methods for nucleosome dissociation: salt or Proteinase K treatment. High salt concentrations promote dissociation by decreasing the attractive force between positively charged histone proteins and negatively charged DNA^13,14^ while Proteinase K is a broad-spectrum serine protease^15^ used for crosslink release and nucleosome digestion in MNase assays^16^. We performed all experiments with the well-characterized yeast strain YJM789, containing approximately 60,000 SNPs relative to the yeast reference genome^17^. In an initial attempt we used 240,000 cells corresponding to approximately 3 ng genomic DNA as input for tagmentation. For comparison, we performed extraction-free preparation with the commercial Nextera enzyme, using buffers and reagents from the kit (Nextera XT, Illumina). As a reference standard we prepared a library from 150 pg extracted YJM789 genomic DNA using reagents from the kit (NXTMP). Paired-end 75-bp reads were generated on a Illumina MiSeq platform and reads mapped to the reference yeast genome (sacCer3 R64.2.1) using *bwa*^18^.

**Figure 1.**
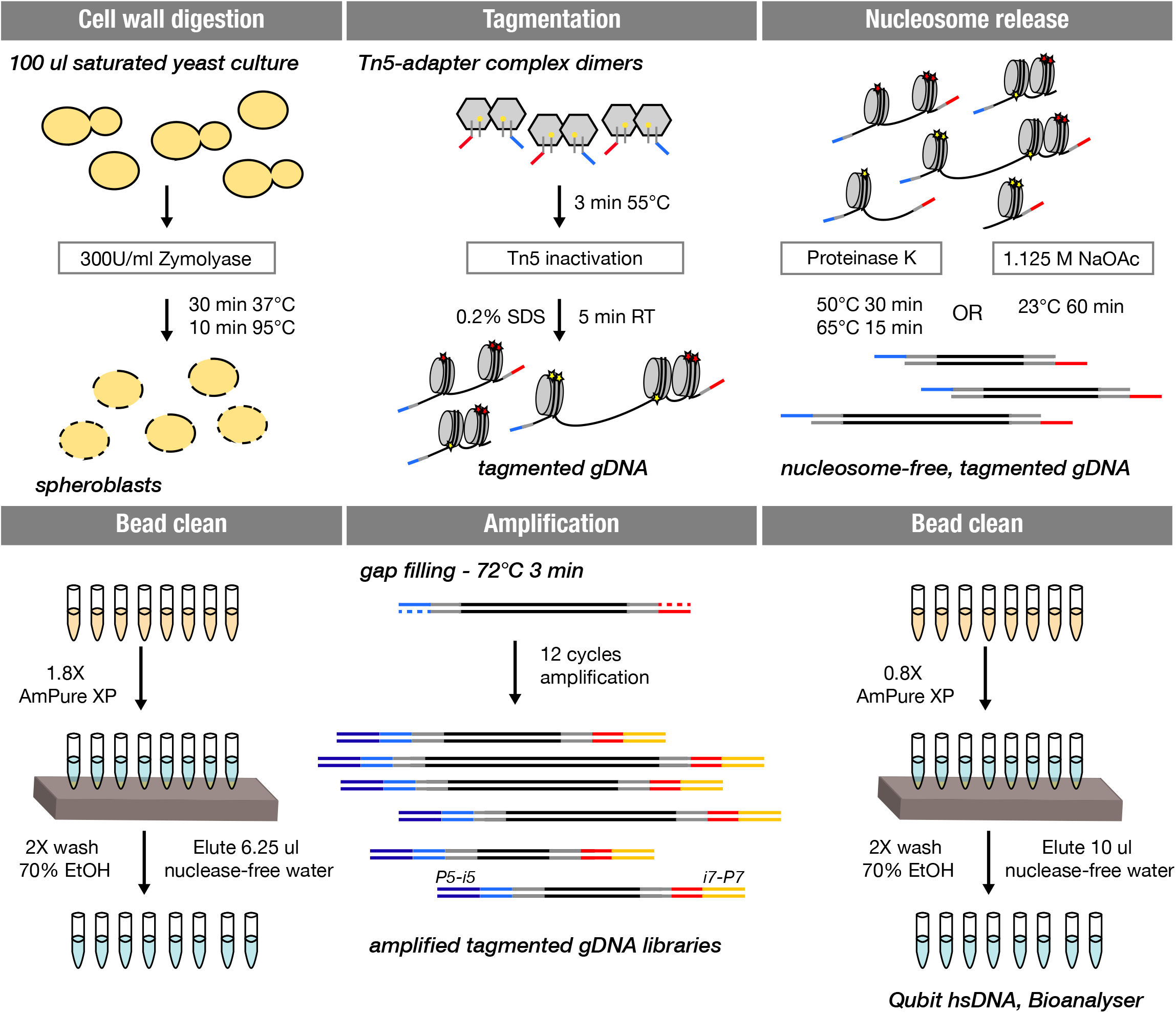
A simplified whole genome sequencing library preparation workflow without genomic DNA isolation. Workflow of cell wall digestion, gDNA tagmentation, nucleosome dissociation and subsequent NGS library preparation for dual index (i5/i7) whole-genome sequencing. The double-stranded part of the linker oligonucleotide is shown in gray with a yellow dot depicting the phosphorylated 3’ end. The 5’ overhangs serving as templates for the indexed P5 (dark blue) or P7 (orange) adapter primers are shown in blue and red, respectively.

A major concern when skipping genomic DNA isolation is the reduced accessibility of genomic DNA to Tn5 and polymerase, leading to a greater variation in coverage across the genome (coverage bias). To assess the impact on the distribution of genomic coverage, we quantified the number of reads mapping to each position, as well as average genome coverage for each sample. We used the ratio of per base coverage to average coverage to illustrate coverage bias - the closer this ratio is to 1, the more evenly the base is covered relative to the rest of the genome. As a secondary bias metric, we calculated the fraction of the genome covered at least 8-fold, reflecting a common variant calling filter. To measure the impact of coverage variation on variant calling, we determined the fraction of true positive single nucleotide polymorphisms (SNPs) and the false positive call rate with increasing sequencing coverage using the extracted sample (NXTMP) as a reference.

With the Nextera enzyme, libraries prepared from cells without nucleosome dissociation (NA) were highly similar to libraries prepared from extracted genomic DNA (Fig. 2, Fig. S1), and a Proteinase K (ProK) nucleosome dissociation step after tagmentation did not provide an additional benefit. Despite equimolar pooling we obtained very low read numbers for the sample with salt treatment and could not evaluate it across the entire range, but at lower sequencing coverage salt treatment seemed to negatively affect variant calling performance (Fig. 2B). With homemade Tn5_R27S,E54K,L372P_ in contrast, both nucleosome dissociation methods improved coverage (Fig. 2A,C, Fig. S1A) and SNP calling rate (Fig. 2B, Fig. S1B) compared to no treatment (NA) when applied after tagmentation. Salt treatment resulted in libraries that were indistinguishable in quality from the extracted sample. Insert size distribution (median/mode) was similar for the extracted sample (NXTMP 115/70) and extraction-free samples with salt treatment, while Proteinase K treatment resulted in larger insert sizes for both Nextera (150/96) and Tn5_R27S,E54K,L372P_ (182/139) (Fig S1A, Table S1). To assess whether providing Tn5 with a nucleosome-free DNA substrate would further improve genome coverage, we applied Proteinase K treatment after Zymolyase incubation but prior to tagmentation. Compared with treatment after tagmentation we observed larger variation in genome coverage and lower performance for other quality metrics (Fig. S2) for both Nextera and Tn5_R27S,E54K,L372P_. The overall worse performance could stem from a higher loss of material during bead-based purification after Proteinase K treatment, as intact genomic DNA might not bind and elute as efficiently as smaller, tagmented genomic DNA, but we did not investigate this further. Overall, these initial experiments showed that high-quality wholegenome sequencing libraries can be generated without a genomic DNA isolation step with both commercial and homemade transposase enzymes.

**Figure 2.**
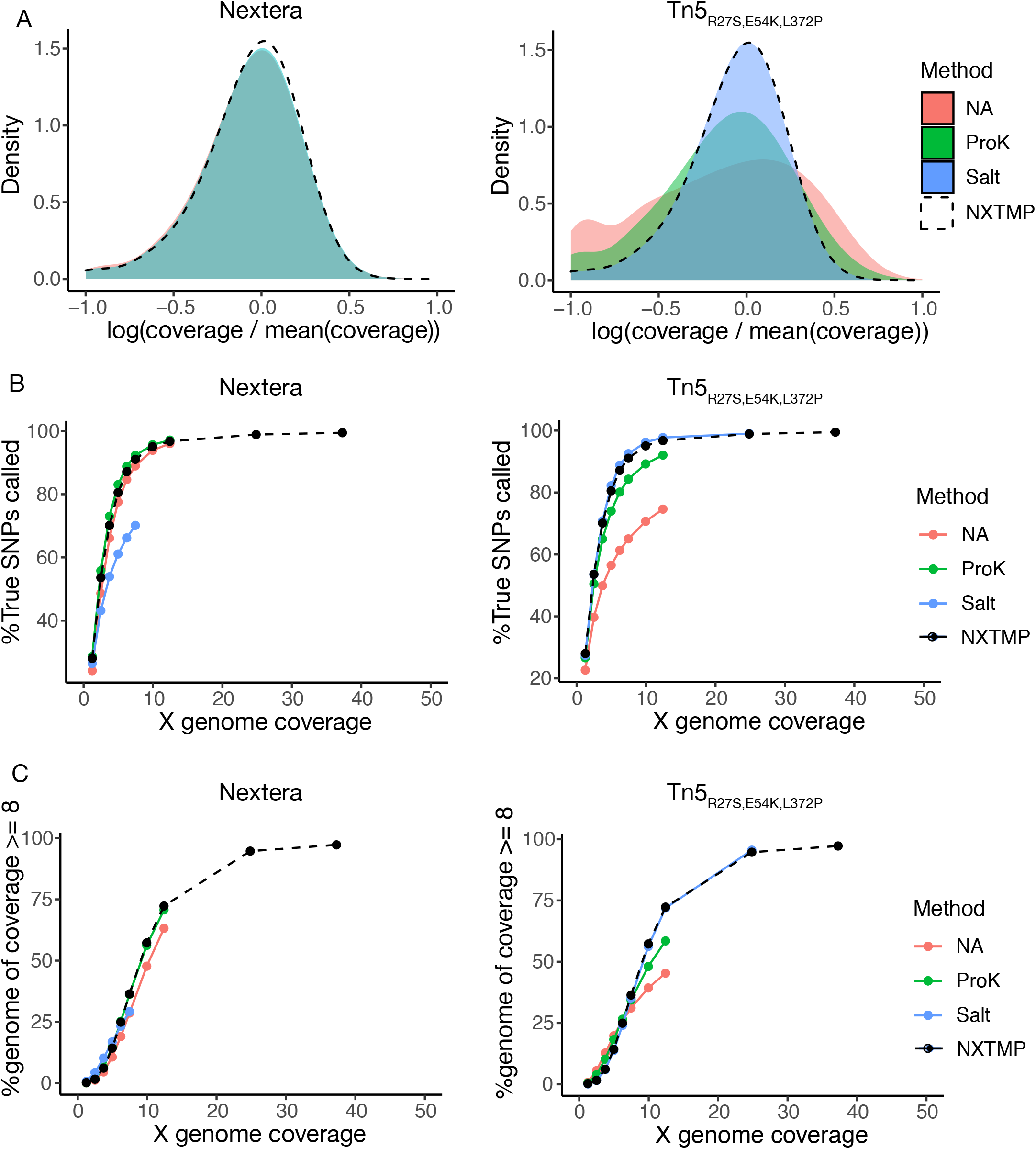
Comparison of different nucleosome dissociation methods in extraction-free library preparation. Samples starting from 240,000 cells were processed following the workflow depicted in Figure 1. After tagmentation nucleosomes were dissociated using salt (blue) or Proteinase K (ProK, green). NA (salmon) indicates the lack of a nucleosome dissociation step. NXTMP (black dashed line) is a library processed from 150 pg extracted genomic DNA with the Nextera XT kit and serves as reference standard. A) Coverage bias distribution (log scale), with bias calculated as coverage at a base divided by average genome coverage. The salt condition is omitted for Nextera due to insufficient reads for this sample. B) Fraction of YJM789 SNPs (in percent) called as a function of sequencing depth (indicated as average genome coverage). The reference set of true positive SNPs (52,373 SNPs) is derived from variant calling on a sample prepared from extracted genomic DNA with the Nextera XT kit (NXTMP). C) Fraction of the genome covered at least 8-fold (in percent) as a function of sequencing depth (indicated as average genome coverage).

### Optimising protocol parameters

Tagmentation tends to generate libraries with shorter insert sizes than methods using mechanical or enzymatic fragmentation^1^, which can be a limitation for applications requiring longer inserts. Compared to Tn5-based standard library preparation from genomic DNA, library preparation from cells with Proteinase K-mediated nucleosome dissociation consistently produced longer fragments, providing an opportunity to address this need. To identify robust conditions for an efficient, extraction-free workflow with our in-house transposase compatible with longer insert sizes we extensively varied different reaction steps. With 65°C at the higher end of the permissible temperature range for Proteinase K activity, we tested incubation at 50°C, and 50°C followed by 15 minutes at 65°C. Both variations improved coverage bias (Fig. S3A) and to a smaller degree variant calling (Fig. S3B, C) compared to incubation at 65°C, but the 50°C treatment by itself led to reduced insert sizes (Fig. S3D, Table S1) and reduced genomic coverage at lower sequencing depths (Fig. S3E). A combined incubation at 50°C followed by 65°C resulted in highest library quality while retaining larger insert sizes.

For initial testing we measured optical density and adjusted cell number of individual cultures, which limits throughput for parallel processing. To evaluate the robustness of the protocol to variable input cell numbers, we processed samples starting from 240,000 to 5 million cells. We adjusted sampling volume from the saturated culture such that 1.25 μl of the Zymolyase reaction contained the desired number of cells, to prevent any changes in the volumes of reactions. Libraries generated from 240,000, 500,000 and 1 million cells (corresponding to approximately 3 ng, 6 ng and 12 ng of input DNA) were highly similar in terms of coverage bias (Fig. 3A, E), variant calling (Fig. 3C, D) and insert size distributions (Fig. 3D). The main effect of increasing cell number was an increased library yield after enrichment PCR. Increasing input cell number to 5 million cells (~60 ng DNA), in contrast, increased coverage bias and resulted in lower SNP-calling rate, potentially due to an unfavorable ratio of Tn5 to DNA molecules or insufficient lysis at higher cell densities. The robustness of the protocol towards cell number variation facilitates parallel processing of many samples as it makes it unnecessary to measure optical density and adjust cell number of individual samples.

**Figure 3.**
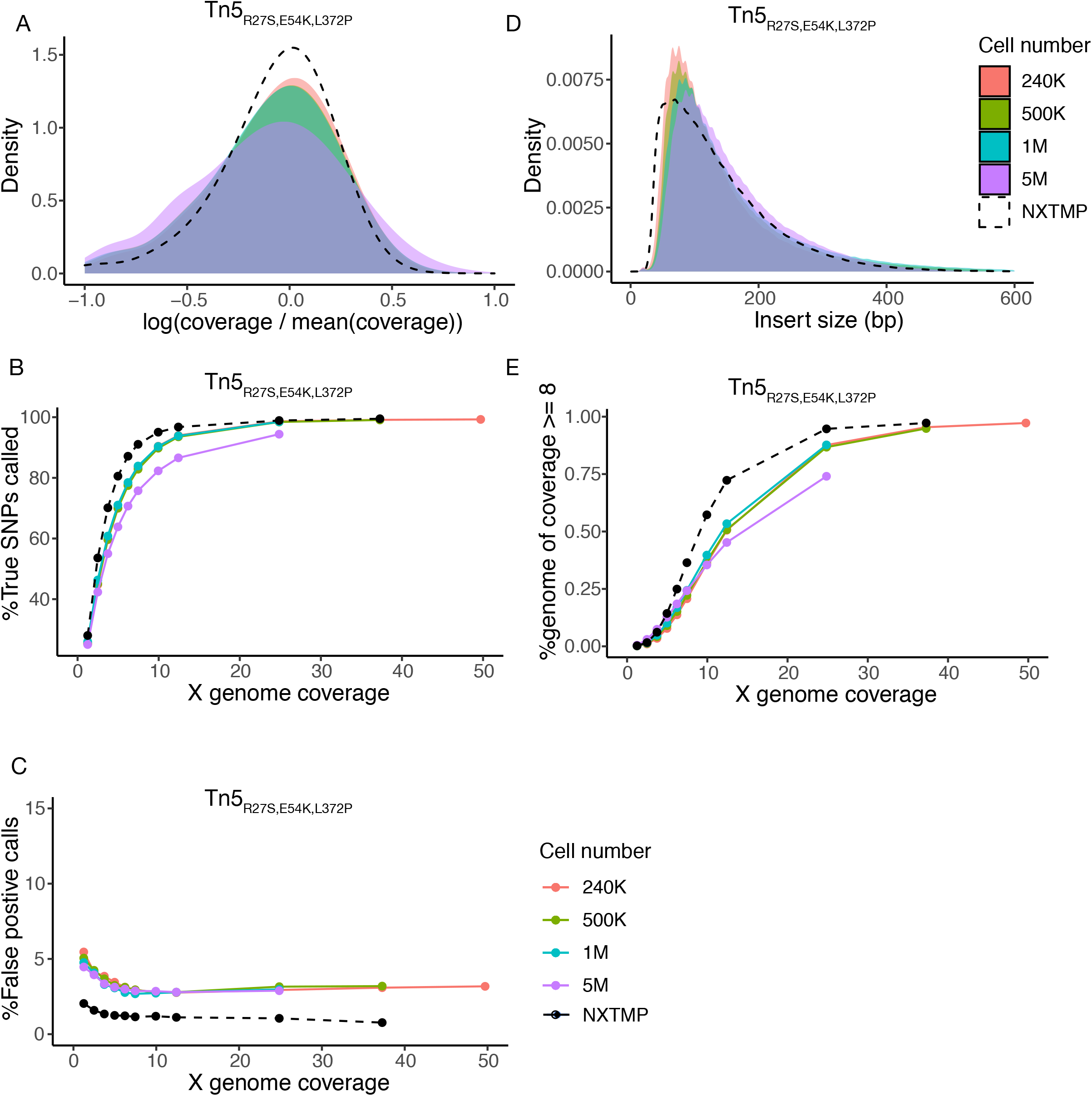
Extraction-free library preparation is robust to variation in input cell number. Samples starting from 240,000 (salmon), 500,000 (green), 1 million (turquoise) or 5 million (purple) cells were processed using homemade Tn5_R27S,E54K,L372P_ and nucleosome dissociation by incubation with Proteinase K after tagmentation. Cell numbers between 240,000 and 1 million yield stable results in terms of library quality. A) Coverage bias distribution (log scale), with bias calculated as coverage at a base divided by average genome coverage. B) Fraction of YJM789 SNPs (in percent) called as a function of sequencing depth (indicated as average genome coverage). The reference set of true positive SNPs (52,373 SNPs) is derived from variant calling on a sample prepared from extracted genomic DNA with the Nextera XT kit (NXTMP). C) Fraction of false positive SNP calls as a function of sequencing depth (indicated as average genome coverage), with false positive call rate = number of mis-called SNPs/total number of SNPs. D) Distribution of fragment insert sizes. E) Fraction of the genome covered at least 8-fold (in percent) as a function of sequencing depth (indicated as average genome coverage). NXTMP (black dashed line) is a library processed from 150 pg extracted genomic DNA with the Nextera XT kit and serves as reference standard.

Previous studies have reported that magnesium chloride and Dimethylformamide (DMF) concentrations in the tagmentation buffer affect performance^3^. As our tagmentation buffer (TB1: 10mM MgCl_2_, 25% DMF and 10mM Tris-HCl, pH 7.6) was optimized for cDNA libraries we varied buffer composition to improve tagmentation of genomic DNA from spheroplasts. We used 500,000 cells in these tests and omitted the nucleosome dissociation step to evaluate the effect of buffer changes only. Lowering the DMF concentration to 20% led to a slight reduction in coverage bias (Fig. S4A, E) and improved variant calling performance (Fig. S4B, C) to almost the same level as the extracted sample. Lowering DMF further to 17.5%, 15% or 10% (only 10% shown in plot) had no clear benefit compared to 20% DMF. In addition to DMF we altered the concentration of both magnesium chloride and Tris-HCl for an optimized tagmentation buffer TB2 containing 8mM MgCl_2_, 20% DMF and 16mM Tris-HCl, pH 7.6 (final concentrations).

As both altered Proteinase K and tagmentation conditions led to small but discernible improvements in library quality individually, we next evaluated them in combination. Taking advantage of the robustness of the protocol to variations in input cell number, we resuspended pellets of 100 μl saturated overnight cultures directly in 25 μl 300U/ml Zymolyase solution, without measuring the optical density of the culture, and used 1.25 μl of this solution, corresponding to approximately 1.2 Mio cells, as input for tagmentation. After tagmentation and inactivation, we incubated each sample with Proteinase K for 30 min at 50°C followed by 15 min at 65°C, and stored samples at −20°C before proceeding with magnetic bead cleanup and enrichment PCR. As our two different homemade Tn5 enzyme versions (Tn5_E54K,L372P_ and Tn5_R27S,E54K,L372P_) had previously shown different characteristics^2^, we compared their performance to address whether one would be superior to the other. Compared with tagmentation in buffer TB1, coverage and variant calling were clearly improved with buffer TB2 for both transposase versions, resulting in libraries indistinguishable in quality parameters from the sample prepared from extracted genomic DNA (Fig. 4 A-C, Fig. S5B) while retaining longer insert sizes (Fig. S5A, Table S1). To assess whether we could further modulate the insert size distribution specifically towards longer inserts, we evaluated alternative tagmentation parameters. We had previously found that higher enzyme dilutions, which decrease the ratio of Tn5 to DNA molecules, resulted in a progressive shift of the insert size distribution towards larger inserts for libraries made from cDNA^2^. We observed a similar effect for libraries prepared with our extraction-free protocol. Using a five-fold higher enzyme dilution we could shift insert sizes from 158/112 (median/mode) to 211/186 (Fig. S6, Table S1). Tn5 transposase is typically used at 55°C in library preparation applications. Lowering tagmentation temperature to 37°C resulted in larger insert sizes (192/149), indicating reduced activity at this temperature, without affecting library quality (Fig. S6). In summary, we identified optimized nucleosome dissociation and tagmentation conditions, as well as strategies to generate libraries with larger insert sizes than is common for Tn5-based adapter insertion. We also demonstrated that our protocol is robust to variation in input cell numbers, which facilitates parallel processing of many samples.

**Figure 4.**
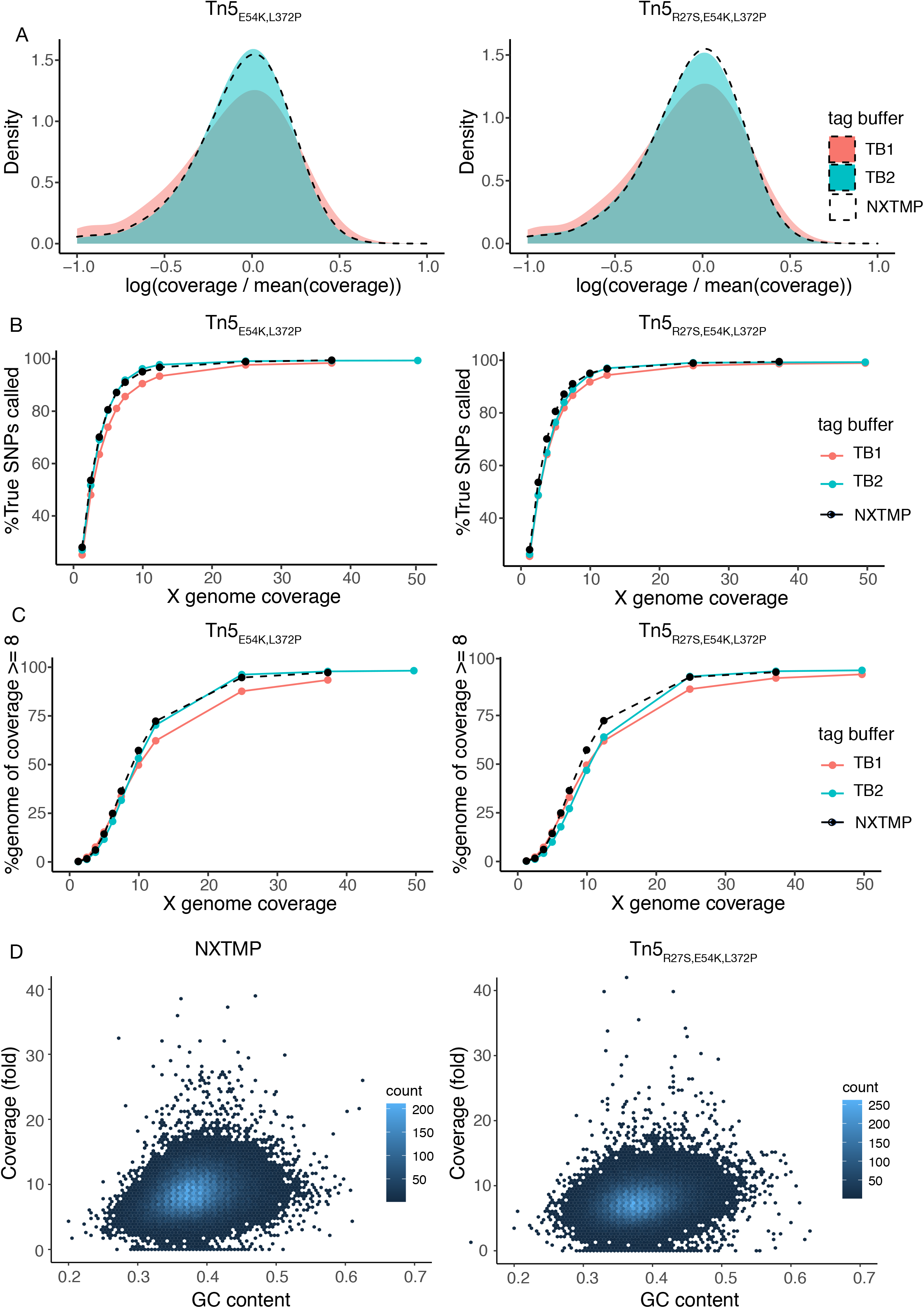
An extraction-free protocol using homemade enzymes produces libraries of the same quality as a commercial, extraction-based solution. Samples were prepared from 100 μl saturated overnight culture with nucleosome dissociation by Proteinase K treatment at 50°C followed by 65°C with two different tagmentation buffers: standard TB1 buffer (salmon) or optimized TB2 buffer (turquoise), and with homemade Tn5_E54K,L372P_ or Tn5_R27S,E54K,L372P_ enzyme. A) Coverage bias distribution (log scale), with bias calculated as coverage at a base divided by average genome coverage. B) Fraction of YJM789 SNPs (in percent) called as a function of sequencing depth (indicated as average genome coverage). The reference set of true positive SNPs (52,373 SNPs) is derived from variant calling on a sample prepared from extracted genomic DNA with the Nextera XT kit (NXTMP). C) Fraction of the genome covered at least 8-fold (in percent) as a function of sequencing depth (indicated as average genome coverage). NXTMP (black dashed line) is a library processed from 150 pg extracted genomic DNA with the Nextera XT kit and serves as reference standard. D) GC content related coverage bias in libraries prepared from extracted genomic DNA with the Nextera XT kit (left panel) or with the extraction-free protocol (TB2 sample using Tn5_R27S,E54K,L372P_ from above, right panel). GC content of the *S. cerevisiae* reference genome was determined for 500 bp sliding windows with 250 bp overlap, and per-base GC content was calculated as the average of overlapping windows.

**Figure 5.**
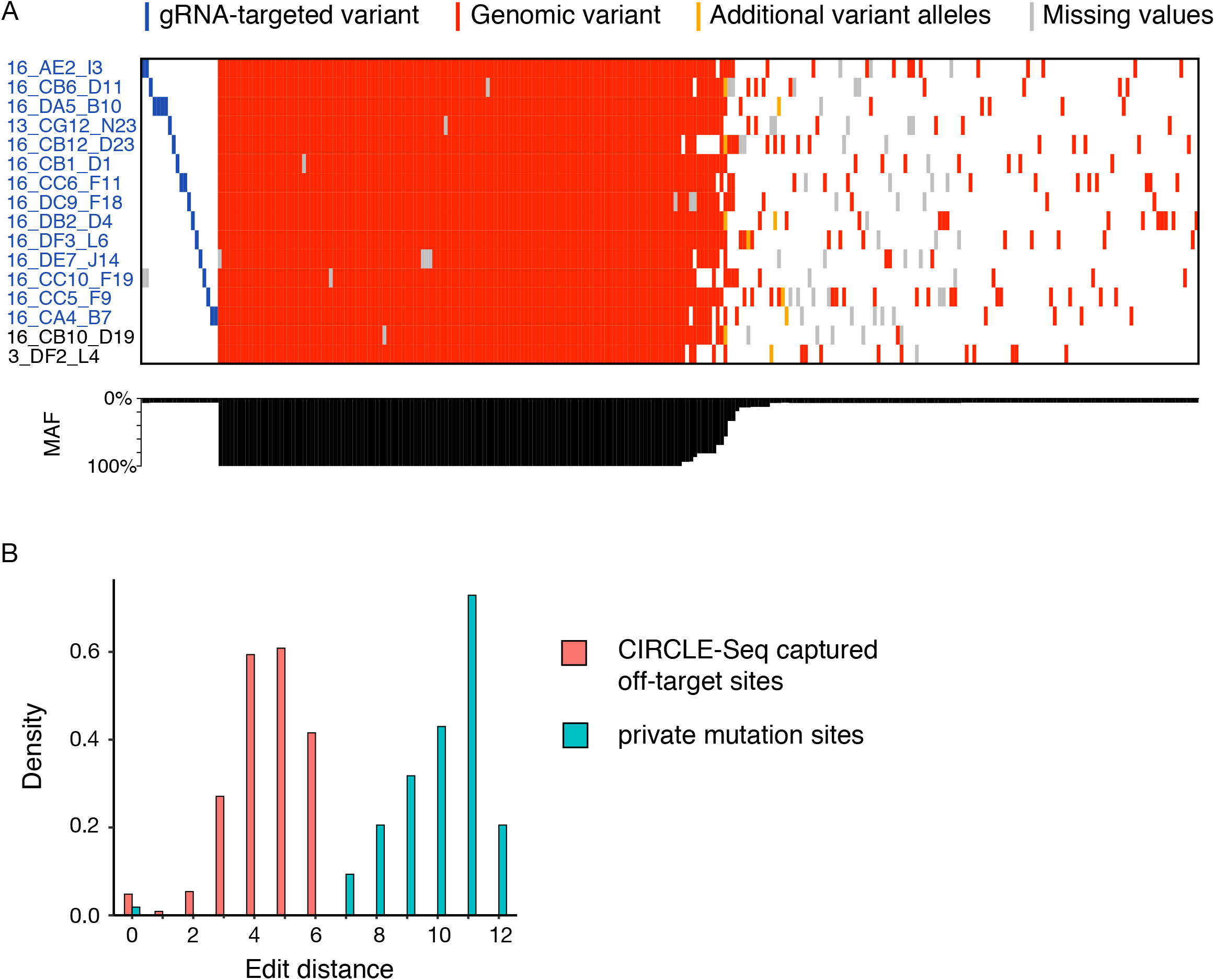
On- and off-target editing analysis in extraction-free libraries. A) Overview of identified mutations in each edited strain (indicated on y-axis), at a sequencing depth of 20X. Designed target variants are shaded in blue. Additional genomic variants are indicated in red or orange, with orange sites representing alternative, non wild-type genotype calls at sites at which an alternative allele has already been detected, likely representing false-positive variant calls. Sites at which no genotype information could be obtained (no coverage) are highlighted in grey. The x-axis depicts a histogram of mutant allele frequency (MAF) of each variant. B) Histogram of edit distances between sequence windows surrounding gRNA targeted variants and alternative variants, reflecting sequence similarity between sites. Sequence similarity between experimentally captured CIRCLE-seq off-target variants (salmon) and their respective gRNA targeted variant indicate expected edit distances if a variant is caused by Cas9-mediated off-target activity. Edit distances obtained for private mutations in each strain in our dataset are depicted in turquoise.

### Detailed characterization of libraries prepared from genomic DNA versus intact cells

We extensively compared the genomic characteristics of libraries generated with the extraction-free protocol, using Tn5_R27S,E54K,L372P_ processed libraries with TB2 as an example, with standard library preparation from extracted genomic DNA (NXTMP). The native Tn5 transposase has an integration site preference^19–21^ and studies have reported a mild GC bias *in vitro*^22^ and in library preparation applications^23^. To assess sequence-dependent bias in coverage we calculated nucleotide composition of the yeast reference genome in 500 bp sliding windows. No correlation with GC content was apparent for either the extracted sample or our extraction-free method (Fig. 4D). The absence of PCR-dependent GC bias, as reported previously^1^, could be a result of using the GC tolerant KAPA HiFi polymerase (KAPA Biosystems).

Coverage of individual positions across the genome was relatively well correlated between standard and extraction-free methods. Positions with high coverage in standard extraction-based library preparation (NXTMP) tended to have high coverage in extraction-free samples, with the notable exception of a population dropping in coverage in the sample without genomic DNA extraction (Fig. S7A). We observed the same trend for extraction-free preparation using Nextera (with Proteinase K treatment, from Fig. 2), indicating the coverage loss was specific to extraction-free protocols and not related to the use of different Tn5 enzymes. We mapped these positions to specific sites in the yeast rDNA locus, which occurs in 100-200 tandem-arrayed 9.1kb repeats on chromosome XII^24,25^, making up almost 10% of the genome. The rDNA locus exists in two distinct chromatin states^26^, an accessible and a highly condensed state, and only a small fraction of the repeats is expressed in normal growth conditions, which could explain the reduced coverage for these regions. This is consistent with a higher variability in coverage across the 37S rRNA gene of the locus specifically in extraction-free samples (Fig. S7B). Across the whole genome, we observed no correlation between coverage and nucleosome positions, as evaluated by ATAC-seq insertion frequency^12^ (Fig. S7C) or nucleosome occupancy^27^ (Fig. S7D), and standard and extraction-free samples again looked very similar. A few outlier positions with distinctly higher coverage in the extraction-free sample, specifically in regions with a nucleosome occupancy score of zero (Fig. S7D), also mapped to the rDNA locus, which shows a bias towards higher coverage of non-transcribed regions (NTS) in extraction-free samples (Fig. S7B). Taken together, we conclude that our extraction-free protocol using homemade Tn5 enzymes generates whole-genome sequencing libraries that are similar in quality to libraries prepared from extracted genomic DNA using a commercial kit, at significantly reduced cost and improved throughput and with the additional benefit of generating longer inserts.

### CRISPR-Cas9 on- and off-target activity profiling in yeast

We applied our method to validate the presence of designed variants and identify any unwanted off-target mutations in a set of 16 yeast strains edited with MAGESTIC, our pooled CRISPR guide-donor method for massively parallel precision editing and genomic barcoding^28^. Unique DNA barcodes present at the genomic barcode locus tag a designed variant in each strain (Table S2). We constructed sequencing libraries of these 16 strains and performed 150PE sequencing, generating an average of ~20-fold coverage per genome. After read mapping, variant calling and variant filtering, we detected 131 single nucleotide polymorphisms and 145 small insertions or deletions across all samples (Fig. 5A). Of the 16 strains, 14 received the desired mutations while two carried the wild-type allele at the targeted locus. Among the 276 called variants, 121 were background variants (mutant allele frequency = 1) representing the baseline genetic differences between the S288c-derivative editing base strain and the S288c reference genome, while 107 were private variants, uniquely present in single lineages (Table S3).

Unintended mutations caused by Cas9 off-target activity should be private mutations, given the uniqueness of each guide RNA sequence and the dependence on sequence similarity to the target^29^. To assess the likelihood of each variant to be derived from an off-target cleavage event, we quantified the sequence similarity of the window surrounding each private mutation to the relevant target sequence (20nt gRNA + 3nt PAM). As a positive control set, we calculated the edit distances between known on- and off-target sequence pairs captured by CIRCLE-Seq^30^ and compared them to the spectrum of edit distances in our dataset. There was a clear difference between the two datasets, with experimentally captured off-target sites from CIRCLE-seq exhibiting edit distances of 0 to 6 to the target site, while all but one of our private variants showed edit distances of 7 or higher (Fig. 5B). Higher edit distances in the CIRCLE-Seq data, corresponding to more mismatches in the off-target relative to the on-target site, were associated with lower cleavage activity (Fig. S8). This further supports that the observed private mutations were unlikely to be caused by Cas9 off-target activity, but rather derived from spontaneous mutation events, accumulated in the cell divisions since editing. In summary, we used our protocol to generate wholegenome sequencing libraries directly from saturated cultures of CRISPR-edited yeast strains, and detected designed on-target, background and spontaneous mutations in all strains. The fact that the majority of the identified mutations were either common (MAF = 1) or private, even at a moderate sequencing depth of 20X, indicates that our extraction-free method generated high-quality, unbiased whole-genome sequencing libraries with even coverage across the genome while substantially simplifying the library construction process and reducing cost.

## CONCLUSION

We describe a simplified whole-genome sequencing library preparation workflow to generate high-quality, sequencing-ready libraries directly from yeast cultures without genomic DNA isolation. Enzymatic digestion of the yeast cell wall is sufficient for Tn5 to access genomic DNA, and removal of residual nucleosomes prior to enrichment PCR can improve the evenness of genomic coverage, such that resulting libraries are indistinguishable in quality from libraries prepared from extracted genomic DNA with a commercial kit (Nextera XT). With our method, the time from saturated yeast culture to library is reduced to less than three hours, enabling massively scaled sequencing projects by eliminating the time- and labor-intensive genomic DNA isolation step while preserving library quality. The robustness of the protocol towards input cell number variation further facilitates parallel processing of many samples as it makes it unnecessary to measure optical density and adjust sampling volumes of individual cultures. In addition to its simplicity and the benefits in throughput, the protocol is highly affordable with reagent costs of 34 cents per sample when using our homemade Tn5 enzymes^2^. This is an approximately 5-10-fold reduction compared to samples prepared from gDNA extracted using the most simple and scalable kit-based methods. Combining our method with low-coverage (3X) sequencing allows genotyping of segregant panels or determining targeted genome mutagenesis outcomes for only $1 per genome. We believe that these advances will enable massively scaled genome sequencing experiments that have so far been hampered by the time, effort and cost spent on genomic DNA isolation.

Libraries generated by transposase-mediated adapter insertion (tagmentation) tend to have shorter insert size distributions compared with methods using mechanical or enzymatic fragmentation. Our method generates longer inserts than a commercial workflow using genomic DNA without the need for size selection, and we identify additional tagmentation parameters such as temperature and Tn5 to DNA ratio that allow flexible modulation of insert size for custom applications. Insert size distributions compatible with Illumina systems can be obtained even with relatively high dilutions of the Tn5 enzyme, which further reduces cost of the assay.

With our method we reliably detected designed on-target and shared background mutations in a panel of edited yeast strains. We could confidently call genotypes at all target sites and detected the majority of mutations, representing differences due to shared genetic background, in all strains. This indicates that our extraction-free method provides high-quality, unbiased genomic information while substantially simplifying the library construction process and reducing cost. While we have only tested this workflow in yeast it should in principle be transferable to other organisms by adjusting initial lysis conditions to the cell type of choice. Adding a homogenization step prior to lysis could further enable direct library preparation from tissues.

## ACKNOWLEDGEMENTS

We thank the EMBL Protein Expression and Purification facility for production of the inhouse Tn5 enzymes, and the EMBL Genomics Core Facility for discussion and support during protocol development. This work was supported by an Advanced Investigator grant from the European Research Council (ERC) under the European Union’s Horizon 2020 research and innovation programme (AdG-742804 - SystGeneEdit to L.M.S.). S.C.V. was supported by an Advanced Postdoc Mobility Fellowship from the Swiss National Science Foundation (grant number P300PA_177909).

## AUTHOR CONTRIBUTIONS

L.M.S. and S.C.V conceived the approach, designed the study and were responsible for coordination of the study. B.P.H. and N.D. advised the study. S.C.V. and C.S.T. performed experiments. S.L. performed computational analysis. S.C.V. and S.L. wrote the manuscript. All authors read, edited and approved the final manuscript. The authors declare no conflict of interest.

## METHODS

### Yeast strain and media

We used the well characterized yeast strain YJM789^17^, a derivative of a yeast isolated from the lungs of an AIDS patient with pneumonia, in all experiments. YJM789 contains approximately 60,000 SNPs with respect to the *S. cerevisiae* reference genome. For all experiments we inoculated a YPAD culture directly from glycerol stock and grew it to saturation overnight. For experiments with genomic DNA, we extracted genomic DNA using Master Pure genomic DNA extraction Kit (Epicentre) following the manufacturer’s instructions. We confirmed quality by gel and quantified genomic DNA yield using the Qubit high-sensitivity DNA assay.

### Cell lysis and adjustment of cell numbers

For Zymolyase treatment, cells from overnight cultures were pelleted at 1000g for 3 minutes and resuspended in 50 μl of 300U/ml Zymolyase 100-T (AMSBIO) solution, incubated at 37°C for 30 minutes followed by 10 minutes at 95°C to inactivate Zymolyase. 1.25 μl of the solution was used for tagmentation. Final desired cell number was adjusted prior to Zymolyase treatment by measuring OD_600_ of a diluted overnight culture and calculating sampling volume assuming 30 Mio cells per 1ml OD 1 culture (based on value from https://bionumbers.hms.harvard.edu/bionumber.aspx?&id=100986&ver=3). Sampling volume was adjusted such that 1.25 μl of the 50 μl Zymolyase reaction contained the desired number of cells.

### Tn5 adapter complex assembly

Tn5 was expressed and purified as previously described^2^. Tagmentation adapters were annealed as previously described^2^ and thawed on ice. As Tn5 storage buffer (20 mM Tris pH 7.4, 800 mM NaCl, 50% Glycerol) contains relatively high amounts of salts we mixed 2 μl of Tn5 protein (0.5 mg/ml) with 0.5 μl of each annealed adapter (70 μM stock) and 8 μl of 20 mM Tris pH 7.5 for adapter loading. Providing adapters in slight excess to Tn5 favors all Tn5 dimer molecules to be occupied by two adapters, shifting the equilibrium to the fully saturated Tn5-adapter complex. The mixture was incubated at 23°C for 30-60 min at 300 rpm in a thermoshaker. For its use in tagmentation, we further diluted the complex using dilution buffer (10 mM Tris pH 7.5, 150 mM NaCl) to the desired dilution factor.

### Tagmentation with homemade Tn5

Unless indicated otherwise 1.25 μl of lysed cells were mixed with 1.25 μl 1:10 diluted, adapter-loaded Tn5_R27S,E54K,L372P_ and 2.5 μl tagmentation buffer TB1 (10mM MgCl_2_, 25% DMF (v/v), 10mM Tris-HCl final concentrations, adjusted to pH 7.6 using acetic acid). Tagmentation buffer without DMF was prepared as a 2X solution and DMF was added fresh immediately before tagmentation for every experiment. Samples were incubated on a pre-heated thermocycler block for exactly 3 minutes at 55°C after which 1.25 μl 0.2% SDS was added immediately for neutralization, and samples were incubated for 5 min at room temperature. For library amplification we added 1.25 μl each of indexed P5 and P7 primer (10uM), 6.75 μl of 2X KAPA HiFi Ready mix (KAPA Biosystems) and 0.75 μl of DMSO to 6.25 μl of tagmentation reaction. Salt or Proteinase K treated samples were purified using 1.8X volumes of AMPure XP (Beckman Coulter) beads prior to PCR amplification. A gap-filling step followed by 12-cycle enrichment PCR was performed as described previously^2^ and samples were purified using 1.8X volumes AMPure XP beads and eluted in 10 μl nuclease-free water. We quantified library yield using the Qubit high-sensitivity DNA kit and evaluated library quality on an Agilent Bioanalyzer (high-sensitivity DNA assay).

### Tagmentation with Nextera enzyme

Samples processed from extracted genomic DNA with Nextera XT kit (Illumina) were prepared following kit instructions with the following changes: We used 150 pg DNA as input and scaled down reaction volumes such that we used ¼ of indicated reagents per reaction. For cell-based protocols with Nextera enzyme, tagmentation, inactivation and library amplification were performed following kit instructions and using commercial reagents, but only using ¼ of indicated reagents. Briefly, 1.25 μl gDNA (100 pg/μl) or zymolyase-treated cells were mixed with 2.5 μl Tagment DNA Buffer (TD) and 1.25 μl Amplicon Tagment Mix (ATM). Samples were incubated on a pre-heated thermocycler block for exactly 3 minutes at 55°C after which 1.25 μl Neutralize Tagment Buffer (NT) was added and samples were incubated for 5 minutes at room temperature. Salt or Proteinase K treated samples were purified using 1.8X volumes of AMPure XP (Beckman Coulter) beads prior to PCR amplification. For enrichment PCR we added 3.75 μl Nextera PCR Master Mix (NPM) and 1.25 μl each of Index N and Index S primers, and ran the NXT PCR program described in the kit instructions. Amplified libraries were purified using 1.8X volumes AMPure XP beads and eluted in 10 μl nuclease-free water. We quantified library yield using the Qubit high-sensitivity DNA kit and evaluated library quality on an Agilent Bioanalyzer (high-sensitivity DNA assay).

### Nucleosome dissociation

We tested two nucleosome dissociation methods. For salt treatment 6.25 μl of tagmented library was incubated with 3.75 μl of 3M NaOAc (final 1.125 M) at room temperature for 1 hour. For Proteinase K treatment 6.25 μl of tagmented library was incubated with 1.75 μl 0.4mg/ml Proteinase K (diluted from 20mg/ml stock) at 65°C for 30 min after tagmentation, unless indicated otherwise. When we applied nucleosome dissociation before tagmentation we combined cells for several samples using 10 μl of the spheroplast solution as input and eluting in the same volume after bead purification.

### Combined changes

100 μl of a saturated overnight culture (no OD measured) was pelleted and re-suspended in 25 μl 300U/ml Zymolyase solution and incubated at 37°C for 30 minutes, followed by 10 minutes at 95°C. We used 1.25Ul of this solution as input for tagmentation, corresponding to approximately 1.2 Mio cells (assuming 30 Mio cells in 1 ml OD 1 culture and saturation at OD 8). Samples were processed with indicated dilutions of Tn5_E54K,L372P_ or Tn5_R27S,E54K,L372P_ using indicated tagmentation buffers (TB1 or TB2) and tagmentation temperatures (37°C or 55°C). After tagmentation with the corresponding enzyme and inactivation, samples were incubated with Proteinase K for 30 min at 50°C followed by 15 min at 65°C and stored at −20°C before proceeding with magnetic bead cleanup (1.8X) and enrichment PCR. Final purification was done with 0.8X AMPure XP beads. The final protocol is also described in file S1. All samples were pooled and paired-end 150-bp reads were generated on a Illumina NextSeq platform. <u>TB1</u>: 10mM MgCl_2_, 25% DMF (v/v), 10mM Tris-HCl final concentrations, adjusted to pH 7.6 using acetic acid. <u>TB2</u>: 8mM MgCl_2_, 20% DMF (v/v), 16mM Tris-HCl final concentrations, adjusted to pH 7.6 using acetic acid). Tagmentation buffer without DMF was prepared as a 2X solution and DMF was added fresh immediately before tagmentation.

### Library preparation for on-and off-target editing analysis

16 randomly picked MAGESTIC-edited yeast strains were inoculated from glycerol stocks into 100 μl YPAD in a 96-well plate, and grown overnight at 800 rpm at 30°C. Entire cultures were pelleted at 1000g for 3 min pellets were resuspended in 25 μl 300U/ml Zymolyase solution and incubated at 37°C for 30 min, followed by 10 min at 95°C. 1.25 μl of the solution was mixed with 2.5 μl TB2 and 1.25 μl of 1:10 diluted Tn5_R27S,E54K,L372P_, and tagmentation was performed at 55°C for 3 min, followed by inactivation with 1.25 μl 0.2% SDS for 5 min at room temperature. For nucleosome dissociation, samples were incubated with 1.75 μl 0.4 mg/ml Proteinase K for 30 min at 50°C followed by 15 min at 65°C and stored at −20°C before proceeding with magnetic bead cleanup (1.8X) and enrichment PCR. Final purification was done with 0.8X AMPure XP beads. All samples were pooled and paired-end 150-bp reads were generated on a Illumina NextSeq platform.

### Whole-genome sequence analysis

Each whole-genome sequencing dataset was down-sampled to consistent numbers of reads for further comparisons, using bash command “gunzip -c *reads.fastq.gz* | head -n *N*”. For comparisons that involved both samples with 150bp and 75bp reads all reads were trimmed to length 75 prior to downstream processing. 3’ Transposon adapter sequences were trimmed off using cutadapt^31^ and trimmed paired-end reads mapped to the S288c yeast reference genome (sacCer3 R64.2.1) using *bwa*^18^ (v0.7.17). The bam files containing all mapped reads were sorted and PCR duplicates filtered using an embedded version of *picard* toolkits in *gatk* v4.0^32^. Base quality score recalibration (BQSR) was performed using “gatk BaseRecalibrator” to detect and correct systematic errors in the original base accuracy scores. Statistics for assessing mapping qualities were generated using samtools^33^ (v1.9) with its “flagstat” option. Insert size distribution was extracted using the command “gatk CollectInsertSizeMetrics”. Variant calling was performed using “gatk HaplotypeCaller” with the haploid option “-ploidy 1”. Low-quality variant calls were annotated and filtered using “gatk VariantFiltration” with expression “QD < 2.0 ║ FS > 60.0 ║ MQ < 40.0 ║ SOR > 3.0 ║ MQRankSum < −12.5 ║ ReadPosRankSum < −8.0”.

### SNP calling rate assessment

We used the set of variants called in a sample processed with the commercial Nextera pipeline as gold standard (Number of total true SNPs), and determined true and false positive SNP calling rates with each protocol variation. For any given sample *X*, the true positive rate was defined as [Number of correctly called SNPs/Number of total true SNPs], while the false positive rate was defined as [Number of mis-called SNPs/Number of total SNPs in *X*].

### Nucleosome occupancy and GC bias analysis

We used two datasets of yeast nucleosome mapping^12,27^. For tiling-array data, nucleosome occupancy was assessed by the log ratio of probe signals between mono-nucleosomal DNA and total genomic DNA samples. For ATAC-seq data, we calculated transposon insertion frequency per base using *pyatac*^12^ (integrated in *nucleoatac*).

We grouped base positions of the rDNA locus into four categories: 35S rRNA genes, external transcribed spacers (ETS), internal transcribed spacers (ITS) and non-transcribed regions (NTS), according to published gene annotations of the S288c reference genome (http://sgd-archive.yeastgenome.org/sequence/S288C_reference/genome_releases/S288C_reference_genome_R64-2-1_20150113.tgz).

GC-content was calculated for the yeast autosomal genome with a 500-bp sliding window stepped by 250 bp. We took the mean value from overlapped windows to represent GC-content bias for single base position. For correlation tests (coverage vs. nucleosome occupancy, coverage vs. GC-content), we randomly subsampled 100,000 positions to reduce computational complexity.

### Off-target variant analysis

For the analysis of 16 strains edited by MAGESTIC, read mapping and variant calling were performed as described above, except: 1) gvcf files were generated using “gatk HaplotypeCaller -ERC GVCF” before genotyping in cohort mode; and 2) additional filters were applied to exclude variants with low read depth (< 0.1 x *sample average base coverage*) and variants with missing genotypes in >=2 samples. Mutant allele frequency (MAF) was calculated by counting fraction of non-reference alleles after excluding missing genotypes. For each private mutation (unique variants present in single strain), we scanned for gRNA-like sequences in both strand directions of a (60+*X*) bp (*X* equals variant length) window centered at variant position. We measured the Levenshtein edit distance between on-target sequence and its best match in the (60+*X*) bp window.

Data of *in vitro* on- and off-target sequence pairs were downloaded from the CIRCLE-seq publication^30^ and edit distance between each pair was calculated accordingly.

## DATA AVAILABILITY

The pETM11-Sumo3-Tn5 plasmids carrying either the Tn5_E54K,L372P_ or Tn5_R27S,E54K,L372P_ construct are available upon request. The sequencing data discussed in this manuscript have been deposited on SRA, accession number xxx.

## FIGURE LEGENDS

**Figure S1.**
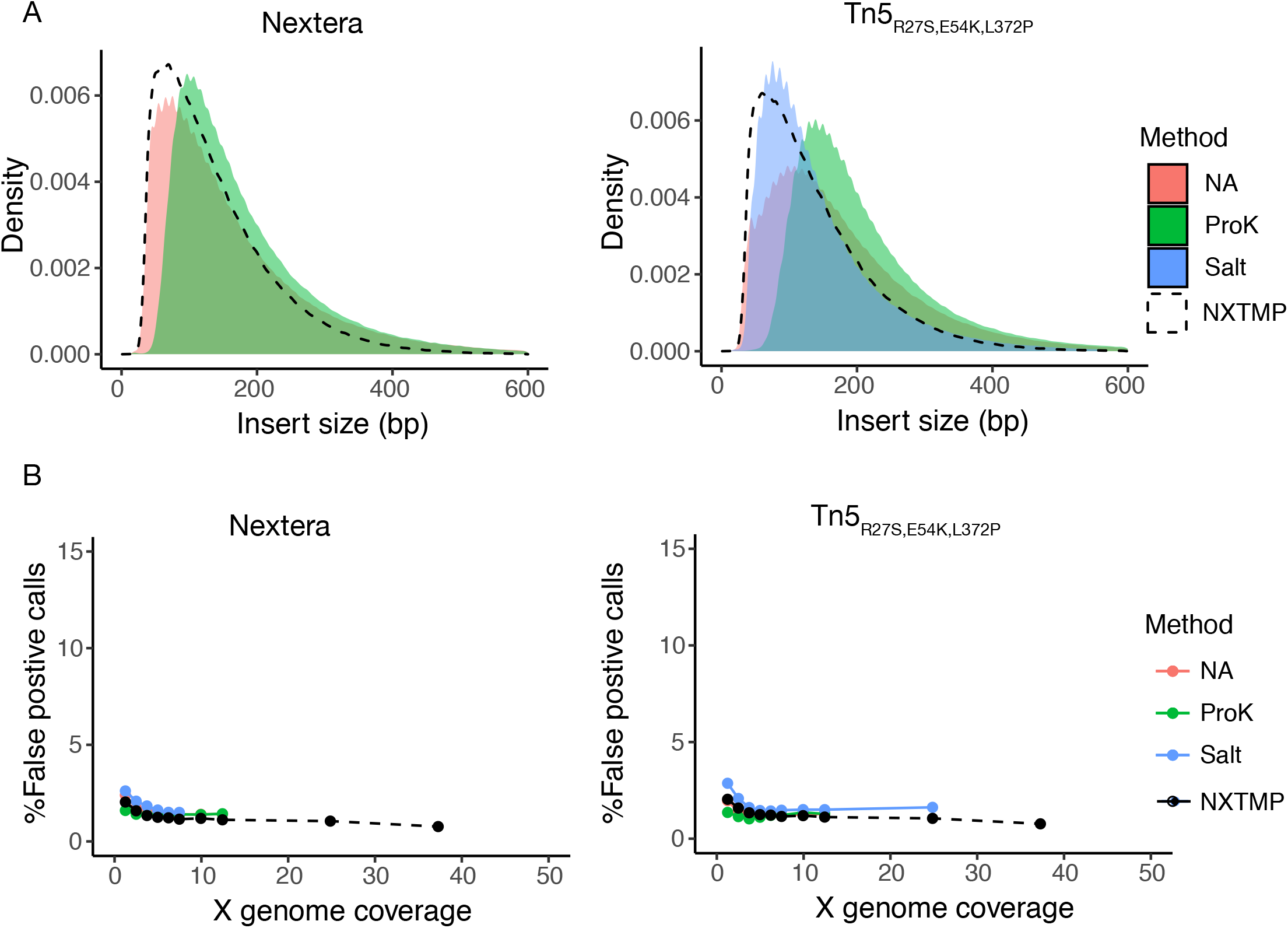
Insert size and false positive calling-rate with different nucleosome dissociation methods in extraction-free library preparation. Salt (blue) indicates nucleosome dissociation by incubation with 1.125 M NaOAc after tagmentation; ProK (green) indicates nucleosome dissociation by incubation with Proteinase K at 65°C after tagmentation; NA (salmon) indicates no nucleosome dissociation step. NXTMP (black dashed line) is a library processed from 150 pg extracted genomic DNA with the Nextera XT kit and serves as reference standard. A) Distribution of fragment insert sizes. The Salt condition is omitted for Nextera due to insufficient reads. B) Fraction of false positive SNP calls as a function of sequencing depth (indicated as average genome coverage), with false positive call rate = number of mis-called SNPs/total number of SNPs.

**Figure S2.**
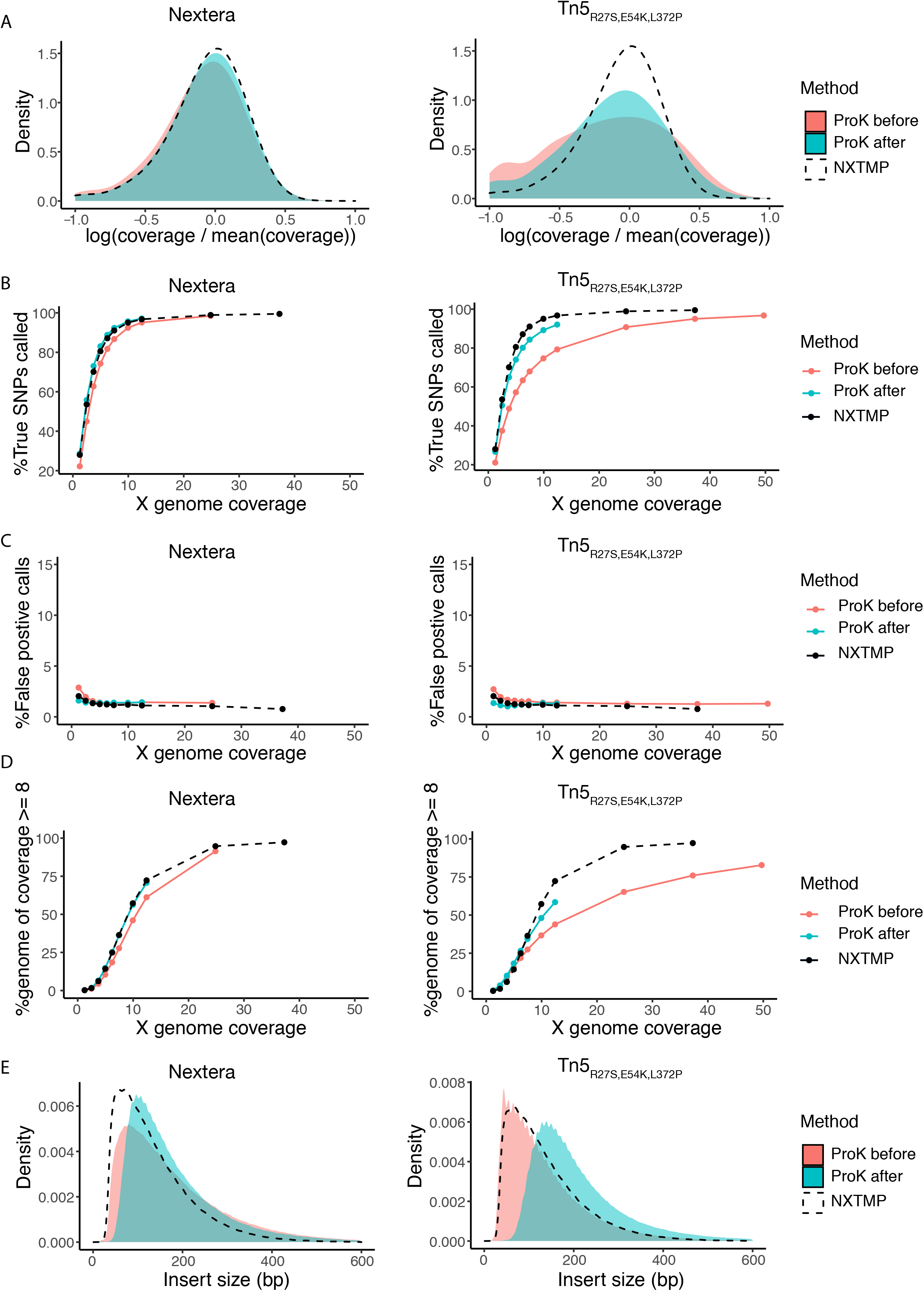
Nucleosome dissociation after tagmentation yields better library quality than pre-tagmentation. Samples starting from 240,000 cells were processed using commercial Nextera enzyme and buffers or homemade Tn5_R27S,E54K,L372P_ enzyme. Nucleosome dissociation by incubation with Proteinase K at 65°C was performed before (salmon) or after (turquoise) tagmentation. A) Coverage bias distribution (log scale), with bias calculated as coverage at a base divided by average genome coverage. The salt condition is omitted for Nextera due to insufficient reads for this sample. B) Fraction of YJM789 SNPs (in percent) called as a function of sequencing depth (indicated as average genome coverage). The reference set of true positive SNPs (52,373 SNPs) is derived from variant calling on a sample prepared from extracted genomic DNA with the Nextera XT kit (NXTMP), and additionally referenced against previously published YJM789 polymorphisms. C) Fraction of false positive SNP calls as a function of sequencing depth (indicated as average genome coverage), with false positive call rate = number of mis-called SNPs/total number of SNPs. D) Fraction of the genome covered at least 8-fold (in percent) as a function of sequencing depth (indicated as average genome coverage). E) Distribution of fragment insert sizes. NXTMP (black dashed line) is a library processed from 150 pg extracted genomic DNA with the Nextera XT kit and serves as reference standard.

**Figure S3.**
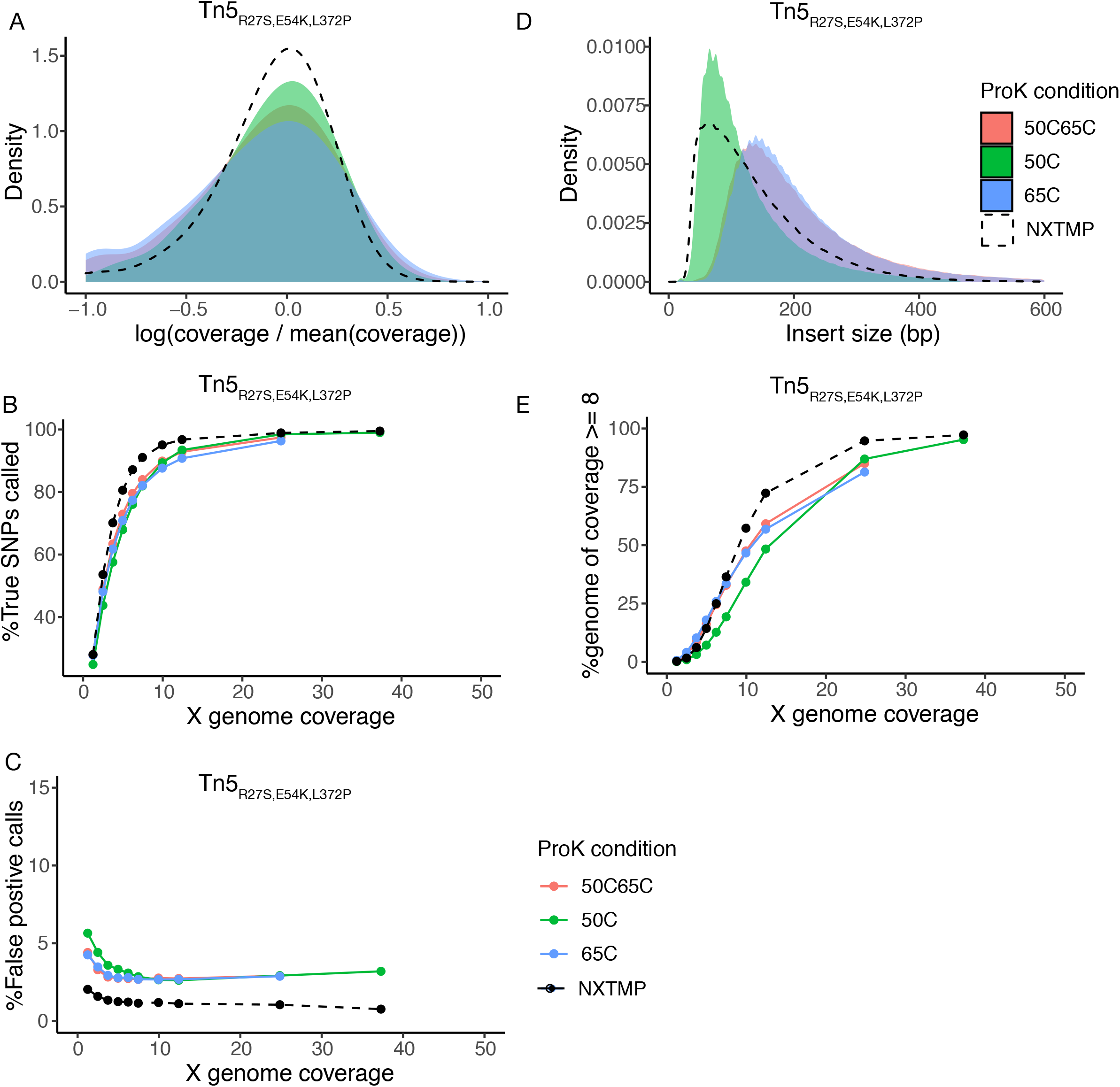
Nucleosome dissociation with variable Proteinase K incubation conditions. Samples starting from 240,000 cells were processed using homemade Tn5_R27S,E54K,L372P_ enzyme and included nucleosome dissociation by incubation with Proteinase K (after tagmentation) at 65°C (blue), 50°C (green) or 50°C followed by 65°C (salmon). A) Coverage bias distribution (log scale), with bias calculated as coverage at a base divided by average genome coverage. The salt condition is omitted for Nextera due to insufficient reads for this sample. B) Fraction of YJM789 SNPs (in percent) called as a function of sequencing depth (indicated as average genome coverage). The reference set of true positive SNPs (52,373 SNPs) is derived from variant calling on a sample prepared from extracted genomic DNA with the Nextera XT kit (NXTMP), and additionally referenced against previously published YJM789 polymorphisms. C) Fraction of false positive SNP calls as a function of sequencing depth (indicated as average genome coverage), with false positive call rate = number of miscalled SNPs/total number of SNPs. D) Distribution of fragment insert sizes. E) Fraction of the genome covered at least 8-fold (in percent) as a function of sequencing depth (indicated as average genome coverage). NXTMP (black dashed line) is a library processed from 150 pg extracted genomic DNA with the Nextera XT kit and serves as reference standard.

**Figure S4.**
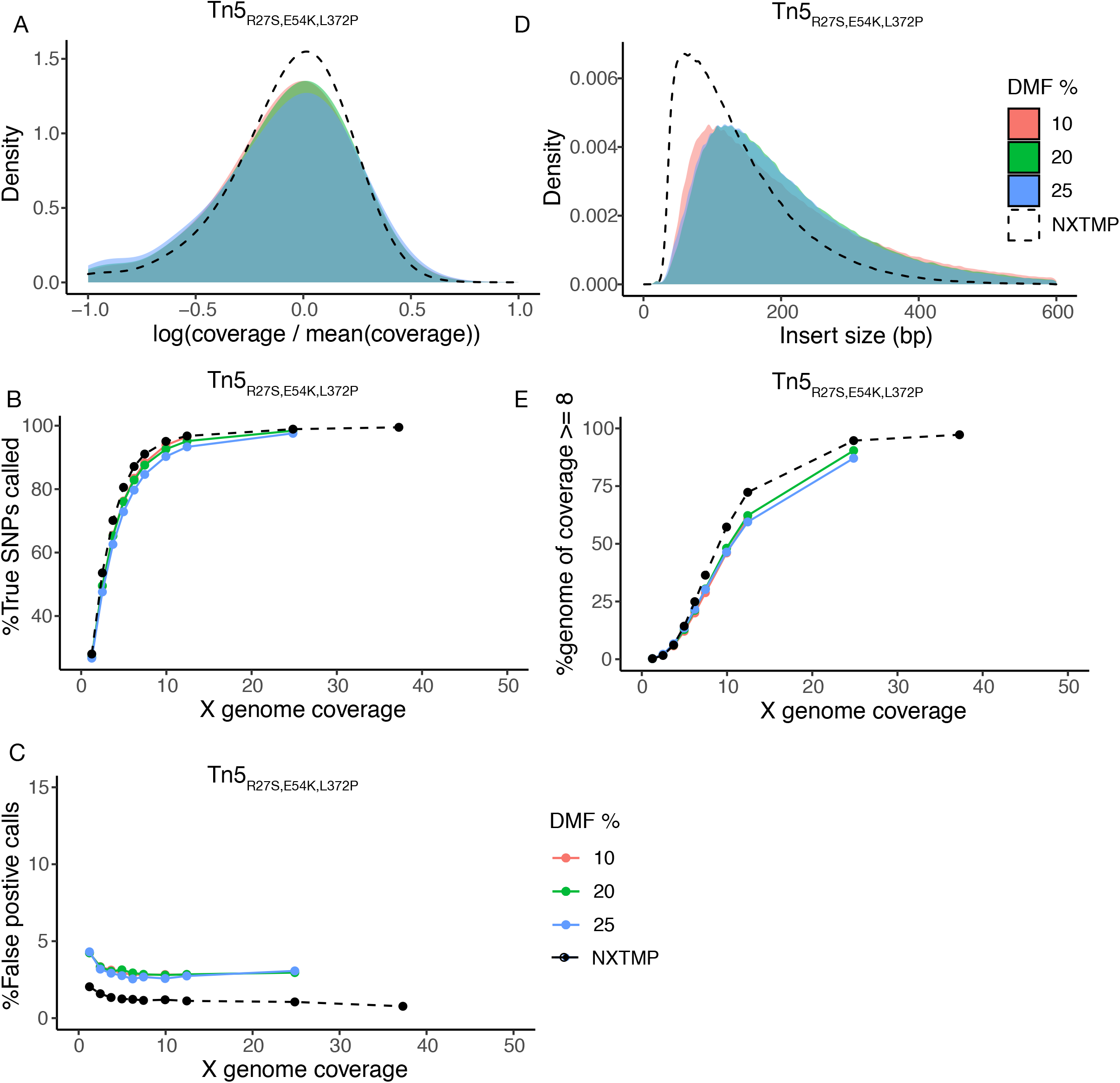
Lower DMF content in the tagmentation buffer improves coverage and variant calling. Samples starting from 500,000 cells were processed using homemade Tn5_R27S,E54K,L372P_ enzyme without a nucleosome dissociation step and with varying DMF percentage in the tagmentation buffer: 25% (blue), 20% (green) or 10% (salmon). A) Coverage bias distribution (log scale), with bias calculated as coverage at a base divided by average genome coverage. The salt condition is omitted for Nextera due to insufficient reads for this sample. B) Fraction of YJM789 SNPs (in percent) called as a function of sequencing depth (indicated as average genome coverage). The reference set of true positive SNPs (52,373 SNPs) is derived from variant calling on a sample prepared from extracted genomic DNA with the Nextera XT kit (NXTMP), and additionally referenced against previously published YJM789 polymorphisms. C) Fraction of false positive SNP calls as a function of sequencing depth (indicated as average genome coverage), with false positive call rate = number of mis-called SNPs/total number of SNPs. D) Distribution of fragment insert sizes. E) Fraction of the genome covered at least 8-fold (in percent) as a function of sequencing depth (indicated as average genome coverage). NXTMP (black dashed line) is a library processed from 150 pg extracted genomic DNA with the Nextera XT kit and serves as reference standard.

**Figure S5.**
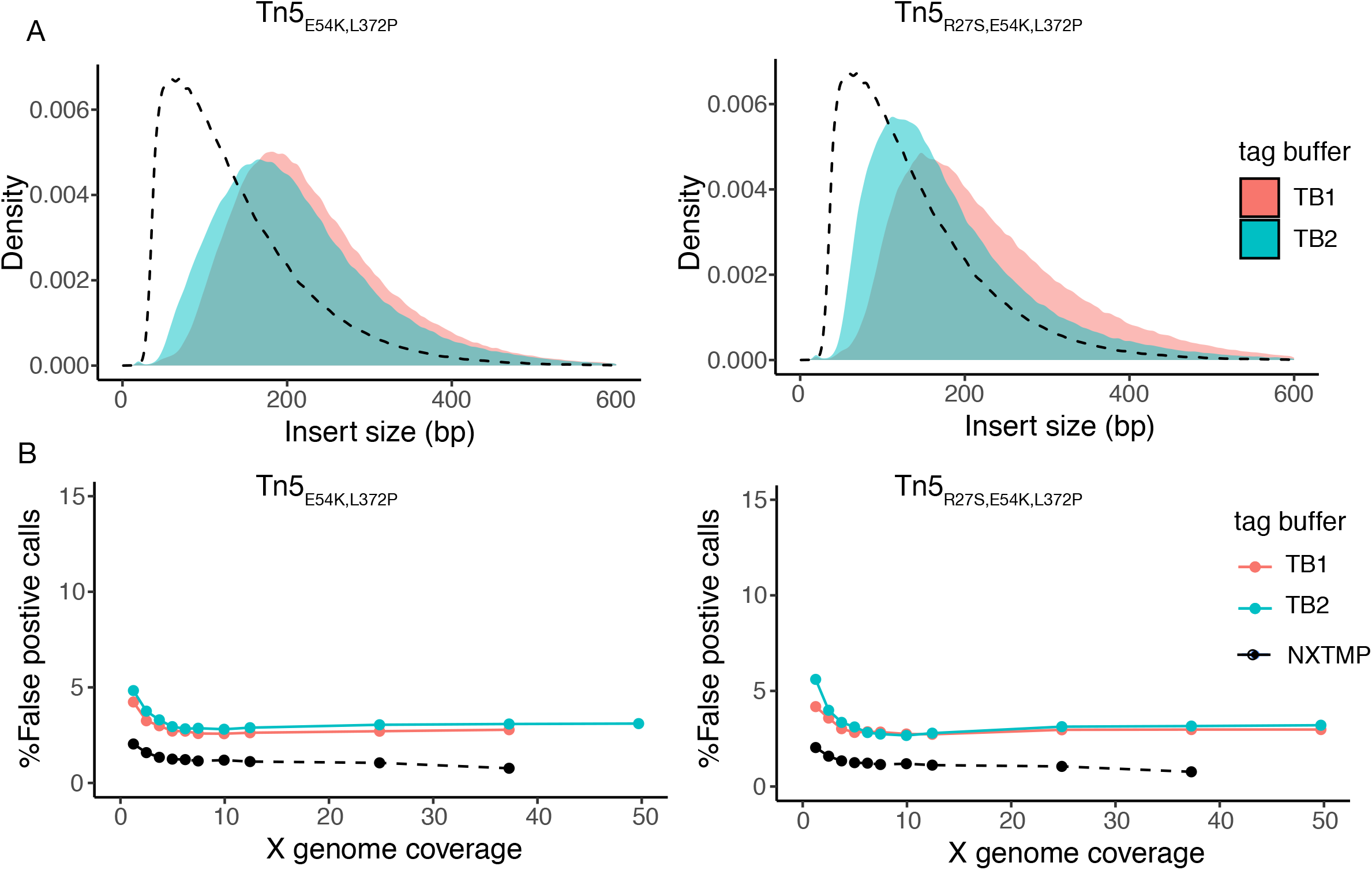
Insert size and false positive calling-rate of libraries prepared with extraction-free protocol using homemade enzymes. Samples were prepared from 100ul saturated overnight culture with nucleosome dissociation by ProK treatment at 50°C followed by 65°C with two different tagmentation buffers, old TB1 buffer (salmon) or optimized TB2 buffer (turquoise), and with homemade Tn5_E54K,L372P_ or Tn5_R27S,E54K,L372P_ enzyme. A) Distribution of fragment insert sizes. B) Fraction of false positive SNP calls as a function of sequencing depth (indicated as average genome coverage), with false positive call rate = number of mis-called SNPs/total number of SNPs. NXTMP (black dashed line) is a library processed from 150 pg extracted genomic DNA with the Nextera XT kit and serves as reference standard.

**Figure S6.**
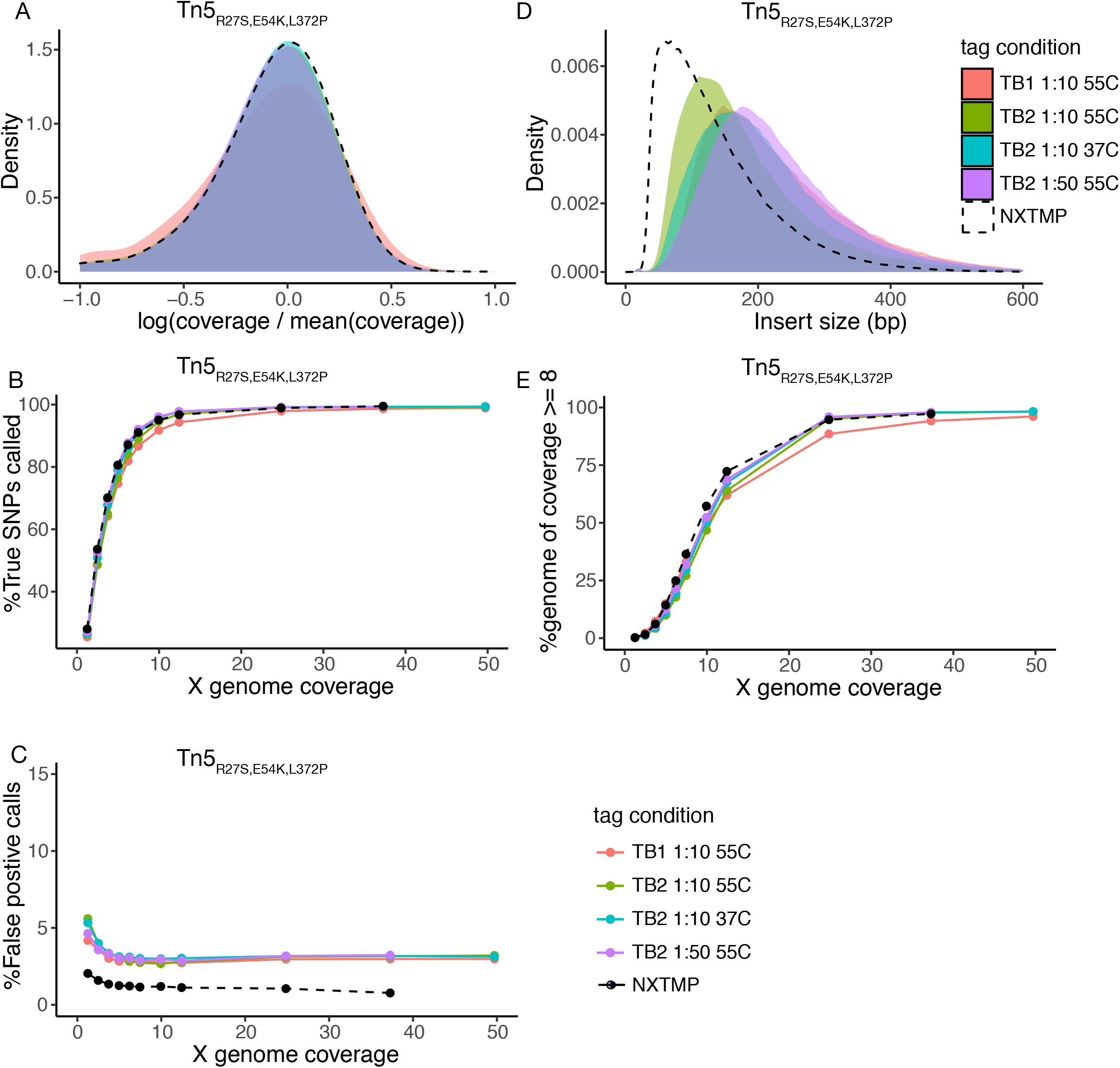
Lower temperature or enzyme concentration during tagmentation increase insert size. Samples were prepared from 100ul saturated overnight culture with nucleosome dissociation by Proteinase K treatment at 50°C followed by 65°C using homemade Tn5_R27S,E54K,L372P_ enzyme and varying tagmentation conditions: tagmentation for 3 min at 55°C in old TB1 buffer, with 1:10 Tn5 dilution (salmon, standard condition); tagmentation for 3 min at 55°C in TB2 buffer, with 1:10 Tn5 dilution (green); tagmentation for 3 min at 37°C in TB2 buffer, with 1:10 Tn5 dilution (turquoise); tagmentation for 3 min at 55°C in TB2 buffer, with 1:50 Tn5 dilution (purple). A) Coverage bias distribution (log scale), with bias calculated as coverage at a base divided by average genome coverage. The salt condition is omitted for Nextera due to insufficient reads for this sample. B) Fraction of YJM789 SNPs (in percent) called as a function of sequencing depth (indicated as average genome coverage). The reference set of true positive SNPs (52,373 SNPs) is derived from variant calling on a sample prepared from extracted genomic DNA with the Nextera XT kit (NXTMP), and additionally referenced against previously published YJM789 polymorphisms. C) Fraction of false positive SNP calls as a function of sequencing depth (indicated as average genome coverage), with false positive call rate = number of mis-called SNPs/total number of SNPs. D) Distribution of fragment insert sizes. E) Fraction of the genome covered at least 8-fold (in percent) as a function of sequencing depth (indicated as average genome coverage). NXTMP (black dashed line) is a library processed from 150 pg extracted genomic DNA with the Nextera XT kit and serves as reference standard.

**Figure S7.**
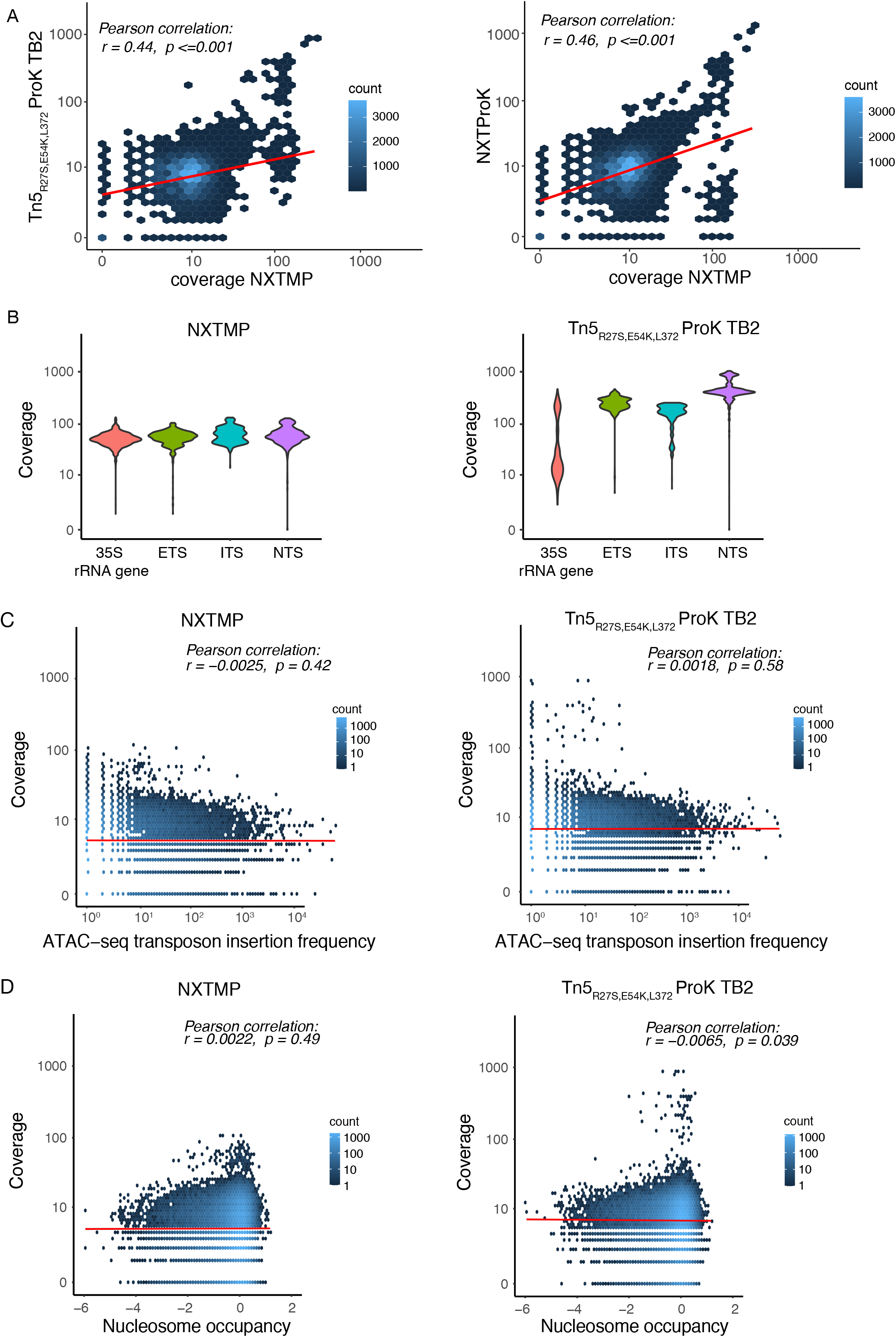
No correlation between genome-wide coverage and nucleosome occupancy in extraction-free libraries. A) Left panel: Correlation of coverage at individual bases between standard (NXTMP) and extraction-free library preparation using homemade enzymes (Tn5_R27S,E54K,L372P_ Prok TB2). Right panel: Correlation of coverage at individual bases between standard (NXTMP) and extraction-free library preparation using Nextera enzyme and commercial reagents. B) Coverage across sequence elements at the yeast rDNA locus in samples with standard (left panel) or extraction-free (right panel) library preparation. C) Correlation between per-base sequencing coverage and ATAC-seq insertion frequency in samples with standard (left panel) or extraction-free (right panel) library preparation. D) Correlation between per-base sequencing coverage and nucleosome occupancy in samples with standard (left panel) or extraction-free (right panel) library preparation. NXTMP = library prepared with Nextera XT kit from 150 pg genomic DNA. Tn5_R27S,E54K,L372P_ Prok TB2 = library prepared from 100ul saturated overnight culture with nucleosome dissociation by ProK treatment at 50°C followed by 65°C and tagmentation in TB2 buffer using Tn5_R27S,E54K,L372P_ (From Fig. 4). NXT ProK = library prepared from 240,000 cells with nucleosome dissociation by Proteinase K treatment at 65°C (from Fig. 2).

**Figure S8.**
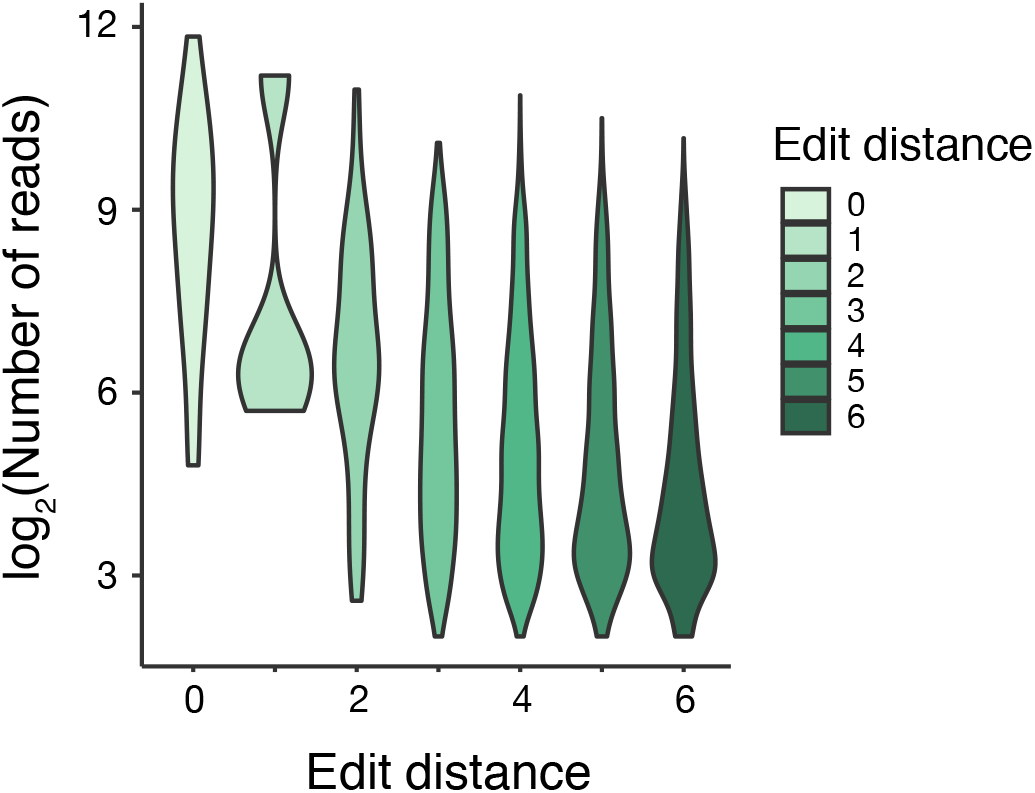
Lower Cas9 cleavage activity at CIRCLE-seq off-target sites with higher edit distances. Violin plots depict distribution of sequencing coverage for each edit distance in the CIRCLE-seq off-target dataset. Due to the nature of the assay, the number of reads across sites in CIRCLE-seq data reflects Cas9-cleavage activity.

**Table S1.** Insert sizes for different tagmentation conditions.

| Sample | Median | Mode | Enzyme | Nucleosome dissociation | Tagmentation buffer | Tagmentation temperature | Enzyme dilution |
| --- | --- | --- | --- | --- | --- | --- | --- |
| NXTMP | 115 | 70 | Nextera | extracted | Nextera | 55°C | none |
| Nextera NA | 129 | 54 | Nextera | NA | Nextera | 55°C | none |
| Nextera ProK | 150 | 96 | Nextera | ProK 65°C | Nextera | 55°C | none |
| Tn5<sub>R27S,E54K,L372P</sub> NA | 151 | 97 | Tn5<sub>R27S,E54K,L372P</sub> | NA | TB1 | 55°C | 1:10 |
| Tn5<sub>R27S,E54K,L372P</sub> ProK | 182 | 139 | Tn5<sub>R27S,E54K,L372P</sub> | ProK 65°C | TB1 | 55°C | 1:10 |
| Tn5<sub>R27S,E54K,L372P</sub> Salt | 118 | 75 | Tn5<sub>R27S,E54K,L372P</sub> |  | TB1 | 55°C | 1:10 |
| Tn5<sub>R27S,E54K,L372P</sub> ProK 50C65C | 178 | 128 | Tn5<sub>R27S,E54K,L372P</sub> | ProK 50°C/65°C | TB1 | 55°C | 1:10 |
| Tn5<sub>R27S,E54K,L372P</sub> ProK 50C | 100 | 65 | Tn5<sub>R27S,E54K,L372P</sub> | ProK 50°C | TB1 | 55°C | 1:10 |
| Tn5<sub>R27S,E54K,L372P</sub> ProK 65C | 180 | 138 | Tn5<sub>R27S,E54K,L372P</sub> | ProK 65°C | TB1 | 55°C | 1:10 |
| Tn5<sub>R27S,E54K,L372P</sub> ProK TB1 | 203 | 145 | Tn5<sub>R27S,E54K,L372P</sub> | ProK 50°C/65°C | TB1 | 55°C | 1:10 |
| Tn5<sub>R27S,E54K,L372P</sub> ProK TB2 | 158 | 112 | Tn5<sub>R27S,E54K,L372P</sub> | ProK 50°C/65°C | TB2 | 55°C | 1:10 |
| Tn5<sub>E54K,L372P</sub> ProK TB1 | 213 | 183 | Tn5<sub>E54K,L372P</sub> | ProK 50°C/65°C | TB1 | 55°C | 1:10 |
| Tn5<sub>E54K,L372P</sub> ProK TB2 | 194 | 167 | Tn5<sub>E54K,L372P</sub> | ProK 50°C/65°C | TB2 | 55°C | 1:10 |
| Tn5<sub>R27S,E54K,L372P</sub> ProK TB2 1:10 37C | 192 | 149 | Tn5<sub>R27S,E54K,L372P</sub> | ProK 50°C/65°C | TB2 | 37°C | 1:10 |
| Tn5<sub>R27S,E54K,L372P</sub> ProK TB2 1:50 55C | 211 | 186 | Tn5<sub>R27S,E54K,L372P</sub> | ProK 50°C/65°C | TB2 | 55°C | 1:50 |

**Table S2.**
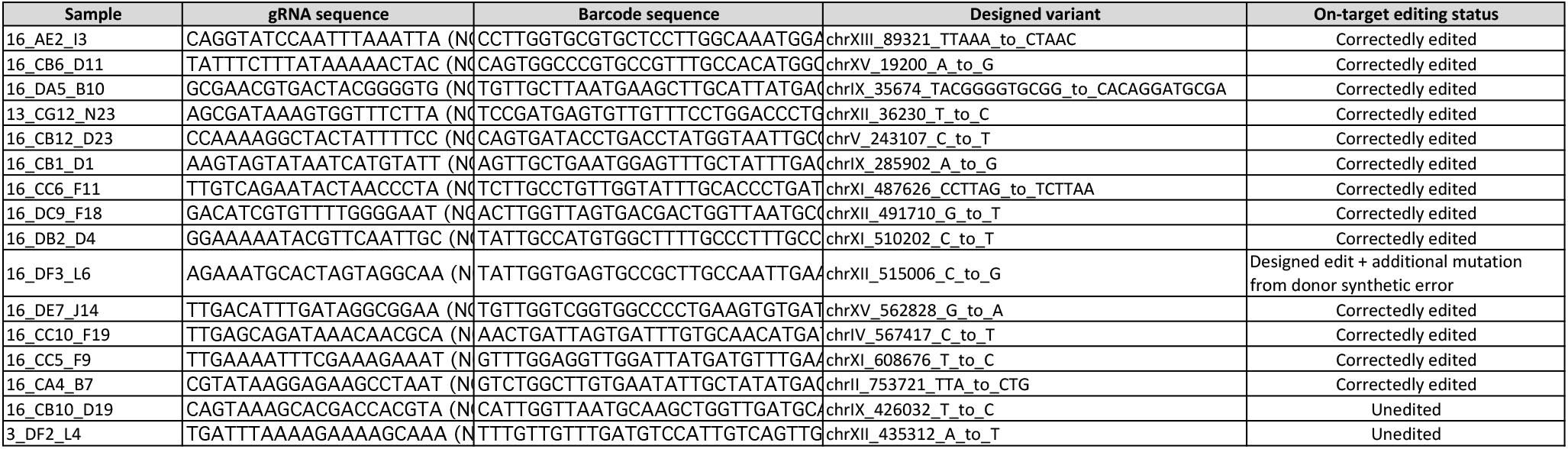
On-target editing outcomes for MAGESTIC-edited CRISPR strains.

**Table S3.**
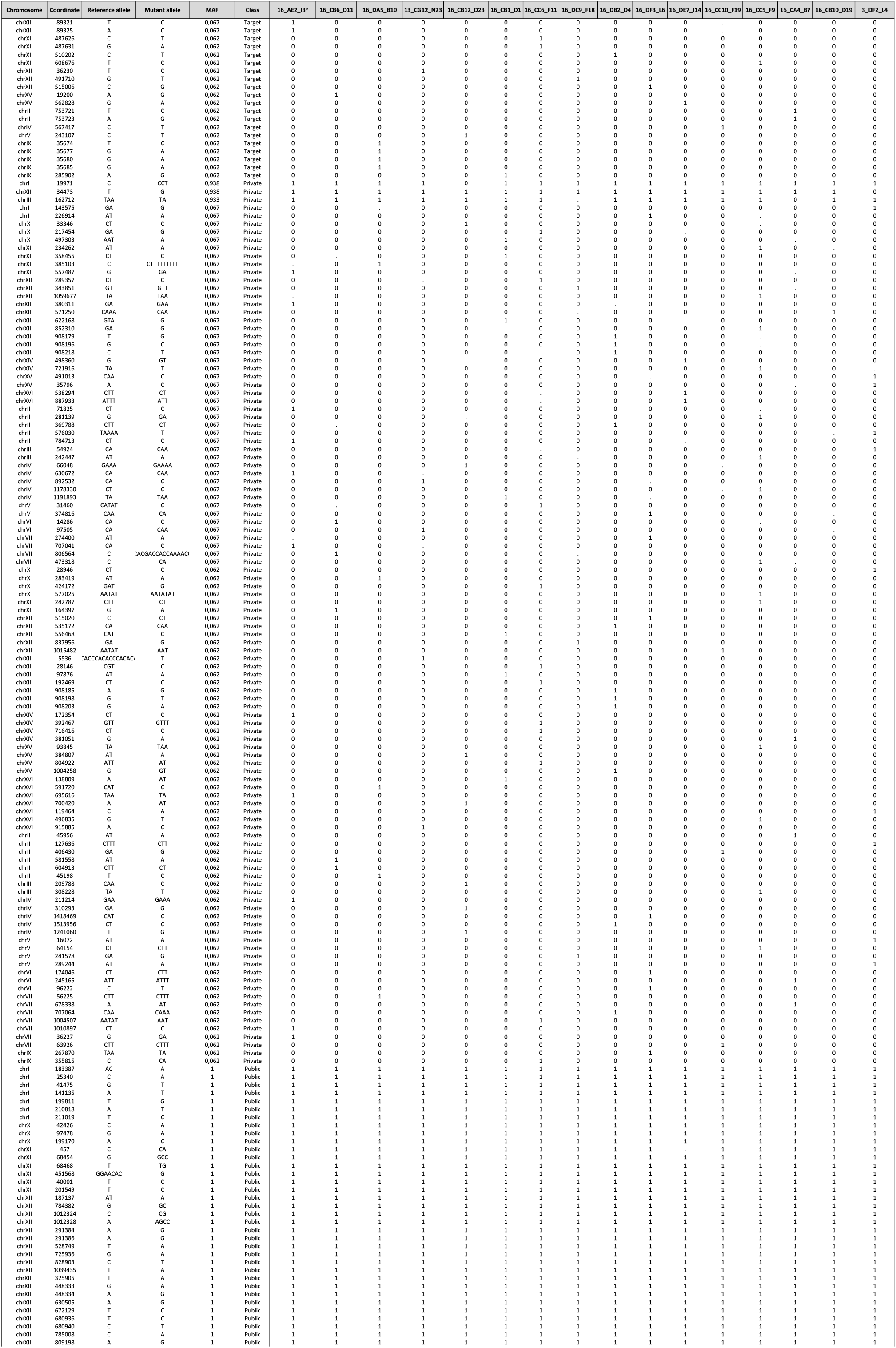
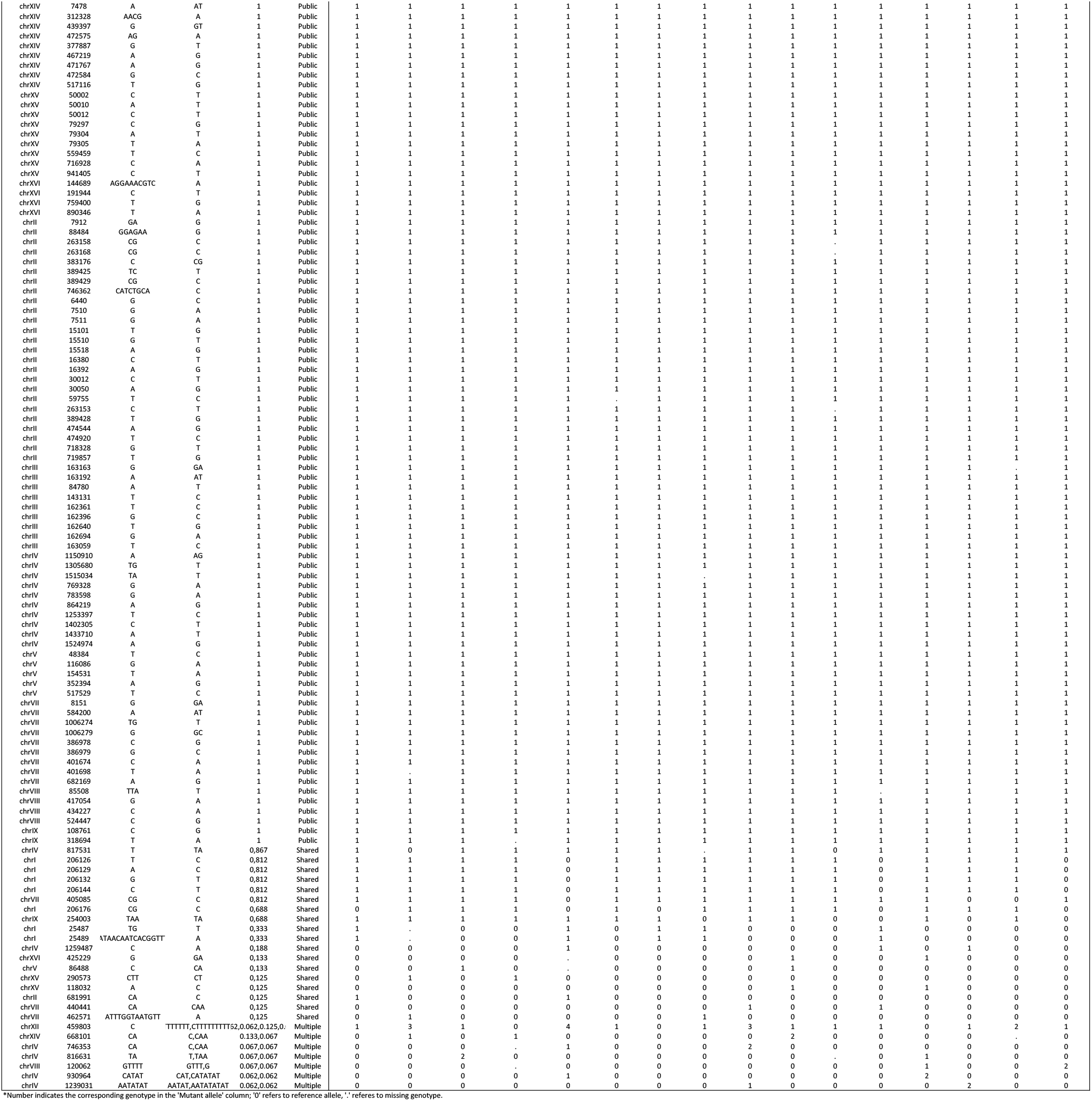
Strain genotypes at polymorphic sites.

## FILE S1. Procedure

**Step 1 - Annealing of the linker.**

*Oligo sequences (from Picelli et al. 2014):*

FC121-1030 5’-TCGTCGGCAGCGTCAGATGTGTATAAGAGACAG-3’
FC121-1031 5’-GTCTCGTGGGCTCGGAGA TGTGTATAAGAGACAG-3’

Tn5MERev 5’-[phos]CTGTCTCTTATACACATCT-3’

1. Resuspend lyophilized oligos at 100uM in annealing buffer
2. Combine fwd and rev linkers 1:1 (FC121-1030 + Tn5MERev and FC121-1031 + Tn5MERev)
3. Anneal in thermocycler:

| Cycle step | Temperature (°C) | Time |
| --- | --- | --- |
| Denaturation | 95 | 5 min |
| Cooling ramp 1 | 95 down to 65 | 0.1°C/s |
| Incubation | 65 | 5 min |
| Cooling ramp 2 | 65 down to 4 | 0.1°C/s |

*store annealed linker in small aliquots of 5-10ul at −20°C*

**Side note - Tn5 linker complex assembly for optimized activity.**

Example for mixing of Tn5 and linkers:

2.0 μl Tn5 protein (0.5 mg/ml – 9.36 uM)

0.5 μl linker A (1030-Tn5MERev annealed oligo)

0.5 μl linker B (1031-Tn5MERev annealed oligo)

8.0 μl Buffer A

11 μl total volume

*Note: this is already a 5.5x dilution of the Tn5 protein.*

*Final buffer composition (approx.)*

*Storage buffer of Tn5 (20 mM Tris pH 7.4, 800 mM NaCl, 50% Glycerol)*

*Salt and Glycerol were diluted 5.5 times (2 μl in total volume of 11 μl)*

*Final Buffer:*

*20 mM Tris pH 7.4*

*145 mM NaCl*

*9 % Glycerol*

*This buffer favors the complex formation between the Tn5 and linkers, as high salt concentration (800 mM NaCl) will inhibit interaction between Tn5 and linkers.*

*Further dilutions of the Tn5 are made with **Buffer B**, which is similar in composition to the buffer in which the Tn5-linker complexes are assembled. Final Tn5 dilutions of 1:10 to 1:50 work well. We recommend testing a dilution series for your application of interest to identify the preferred fragment size spectrum.*

**Calculations:**

2.0 μl of Tn5 (0.5 mg/ml) - 9.36 μM = 18.72 pmol

1 μl of linker mix - 50 μM = 50 pmol

**Ratio:**

Tn5:linkers = 1 : 2.67

***Comment:***

*This ratio favors all Tn5 dimer molecules to be occupied by two linkers. Providing a slight excess of the linkers shifts the equilibrium to the fully saturated Tn5-linker complex.*

**Step 2 - Tn5 loading.**

*Thaw annealed linker on ice!*

*Avoid freeze-thaw cycles of Tn5 (aliquot stocks)*

1. Add 0.5 ul of each linker (from above) to 2 ul Tn5 (0.5 mg/ml) stock and 8ul of Buffer A
2. Mix well and incubate at 23°C for 30 – 60 min at 300 rpm (thermomixer)
3. Dilute the Tn5 in Buffer B to the final desired dilution

*Tn5 is ready to use!*

*If ideal dilution for desired fragment size is not known, make different final dilutions of Tn5 (e.g 1:10, 1:20, 1:50) and run tests for fragment size.*

**Step 3 - Cell wall digestion.**

*Use up to 100ul of saturated overnight culture (between 250,000 and ~2Mio cells final works well)*

1. Spin down cells and resuspend in 25 ul 300U/ml Zymolyase (dilute Zymolyase in nuclease-free water)
2. Incubate 30 min at 37°C, inactivate 10 min at 95°C

**Step 4 - Tagmentation.**

**Important: pre-heat block to desired temperature and make sure to exactly adhere to the incubation time (put samples on ice immediately after tagmentation).**

*Note: Tagmentation can be done at 37°C or 55°C. Fragment size spectrum can be adjusted by changing incubation temperature, time and the dilution factor of the loaded Tn5.*

Tagmentation mix:

1.25 ul zymolyased cells
1.25 ul Tn5 dilution
2.5 ul tagmentation buffer + 20% DMF

mix well, spin down and incubate in pre-heated thermocycler

incubate 3 min at 55°C

put sample on ice

add 1.25 ul 0.2% SDS immediately

mix well by vortexing, spin down and incubate 5 min at room temperature

**Step 5 Version A - Nucleosome release via Proteinase K.**

1.75 ul 0.4 mg/ml Proteinase K

6.25 ul tagmented cells

incubate 30 min at 50°C, 15 min at 65°C

*optional pause point: protocol can be stopped here and samples frozen at −20C*

**Step 5 Version B - Nucleosome release via salt.**

3.75 ul 3M NaOAc

6.25 ul tagmented cells

incubate 60 min at room temperature.

**Step 6 - 1.8X AMPure beads cleanup** (5 min incubation of sample with beads at room temperature, two washes with 70% ethanol), elute in 6.25 ul nuclease-free water

**Step 7 - PCR.**

The oligos used in the PCR contain the sequencing adapter (grafting primer and index barcode)

|  |  |  |
| --- | --- | --- |
| 72°C | 3 min |  |
| 95°C | 30 s |  |
| 95°C | 10 s |  |
| 55°C | 30 s | x 12 cycles |
| 72°C | 60 s |  |
| 72°C | 5 min |  |
| 10°C | hold |  |

**Step 8 - clean up** with 0.8x AmPure beads (to filter out small fragments), or 1.8x (recover everything), elute in 10 ul EB or nuclease-free water

**Step 9 - QC** dsDNA HS Qubit measurement and Bioanalyzer (Qubit yield should be minimum 1 ng/ul when starting from 100 ul saturated culture)

**<u>Buffers:</u>**

**Annealing buffer:**

40mM Tris pH 8

50mM NaCl

**Buffer A:**

20 mM Tris pH 7.5

**Buffer B:**

20 mM Tris pH 7.5

150 mM NaCl

**Tagmentation buffer 2 (TB2, 100ml, in H_2_0, store at room temperature):**

2ml 1M Tris (final 16 mM)

1ml 1M MgCl_2_ (final 8 mM)

adjust pH to 7.6 with 100% acetic acid

Always fresh: add 20% (v/v) DMF

